# Synaptic Transmission Parallels Neuromodulation in a Central Food-Intake Circuit

**DOI:** 10.1101/044990

**Authors:** Philipp Schlegel, Michael J. Texada, Anton Miroschnikow, Andreas Schoofs, Sebastian Hückesfeld, Marc Peters, Casey M. Schneider-Mizell, Haluk Lacin, Feng Li, Richard D. Fetter, James W. Truman, Albert Cardona, Michael J. Pankratz

## Abstract

NeuromedinU is a potent regulator of food intake and activity in mammals. In *Drosophila*, neurons producing the homologous neuropeptide hugin regulate feeding and locomotion in a similar manner. Here, we use EM-based reconstruction to generate the entire connectome of hugin-producing neurons in the *Drosophila* larval CNS. We demonstrate that hugin neurons use synaptic transmission in addition to peptidergic neuromodulation and identify acetylcholine as a key transmitter. Hugin neuropeptide and acetylcholine are both necessary for the regulatory effect on feeding. We further show that subtypes of hugin neurons connect chemosensory to endocrine system by combinations of synaptic and peptide-receptor connections. Targets include endocrine neurons producing DH44, a CRH-like peptide, and insulin-like peptides. Homologs of these peptides are likewise downstream of neuromedinU, revealing striking parallels in flies and mammals. We propose that hugin neurons are part of a physiological control system that has been conserved at functional, molecular and network architecture level.

## Introduction

Multiple studies have demonstrated functional conservation of fundamental hormonal systems for metabolic regulation in mammals and *Drosophila.* This includes insulin [1,2], glucagon [3], and leptin [4]. In addition to these predominantly peripherally released peptides there is a range of neuropeptides that are employed within the central nervous systems (CNS) of vertebrates and have homologs in invertebrates, e.g. neuropeptide Y (NPY), corticotropin-releasing hormone (CRH) or oxytocin/vasopressin [5–9]. Among these, neuromedinU (NMU) is known for its profound effects on feeding behavior and activity: NMU inhibits feeding behavior [10,11], promotes physical activity [12,13], and is involved in energy homeostasis [14,15] and stress response [16,17]. Hugin is a member of the pyrokinin/PBAN (pheromone biosynthesis activating neuropeptide) peptide family and a *Drosophila* homolog of NMU that has recently gained traction due to similar effects on behavior in the fly: increased hugin signaling inhibits food intake and promotes locomotion [18–20]. In addition, both NMU and hugin are also found in analogous or even homologous regions of the mammalian*/Drosophila* CNS. In mammals, distribution of the NMU peptide, NMU-expressing cells and NMU-positive fibers is wide and complex. High levels of NMU have been reported in the arcuate nucleus of the hypothalamus, the pituitary, the medulla oblongata of the brain stem, and the spinal cord [10,21–23]. The number of neurons involved and their morphology is unknown. In *Drosophila,* the distribution of hugin is less complex, yet similar: the peptide is produced by neurons in the subesophageal zone (brain stem) that have hugin-positive projections into the ring gland (pituitary gland), the pars intercerebralis (hypothalamus) and ventral nerve cord (spinal cord) [18]. While comparisons across large evolutionary distances are generally difficult, these regions of the fly brain were suggested to correspond to aforementioned regions of NMU occurrence based on morphological, genetic and functional similarities [24,25] (Fig. 1). Consequently, NMU/hugin has previously been referred to as a clear example of evolutionary constancy of peptide function [26]. Although functional and morphological aspects of neurons employing either neuropeptide have been extensively studied in the past, knowledge about their connectivity is fragmentary. While large scale connectomic analyses in vertebrates remain challenging, generation of high resolution connectomes has recently become feasible in *Drosophila* [27–30].

**Figure 1.**
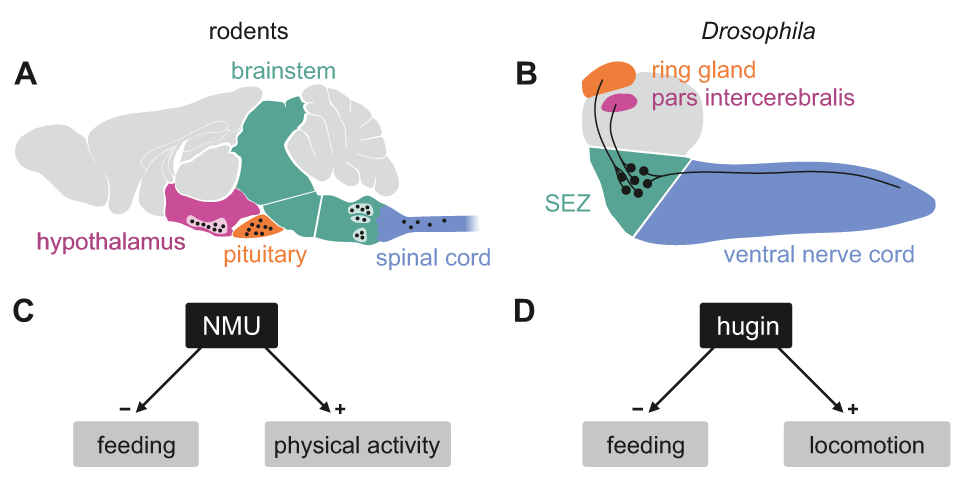
Comparison of mammalian neuromedinU and *Drosophila* hugin. **A,** NeuromedinU (NMU) is widely distributed in the rodent CNS. NMU peptide, NMU-expressing cells and NMU-positive fibers are found in several regions of the brain stem, hypothalamus, pituitary and spinal cord. **B**, In *Drosophila,* distribution of the homologous neuropeptide hugin is less complex but shares similar features: hugin is expressed by sets of neurons in the subesophageal zone (SEZ) that project into the pars intercerebralis, ring gland and ventral nerve cord. Based on morphological, genetic and functional similarities, these regions have been suggested to be analogous and in case of the pars intercerebralis even homologous to regions of the mammalian CNS. Analogous/homologous regions in A and B are given corresponding colors. **C**, **D**, Increased NMU and hugin signaling has similar effects: feeding behavior is decreased whereas physical activity/locomotion is increased.

We took advantage of this and performed an integrated analysis of synaptic and G-protein coupled receptor (GPCR)-mediated connectivity of hugin neurons in the CNS of *Drosophila.* Our data demonstrates that hugin neurons employ small molecule transmitters in addition to the neuropeptide. We identify acetylcholine as a transmitter that is employed by hugin neurons and required for their effect on feeding behavior. Next, we show that hugin neurons form distinct units, demonstrating that clusters of neurons employing the same neuropeptide can be remarkably different in their synaptic connectivity. One unit of hugin neurons is presynaptic to subsets of median neurosecretory cells (mNSCs) in the *Drosophila* homolog of the mammalian hypothalamus. In parallel to the synaptic connectivity, mNSCs also express the G-protein coupled receptor PK2-R1, a hugin receptor, rendering them targets of both fast synaptic transmission and neuromodulatory effects from hugin neurons. These mNSCs produce diuretic hormone 44 (DH44, a CRH-like peptide) and *Drosophila* insulin-like peptides both of which have homologs that are likewise downstream of NMU in mammals [31,32]. Endocrine function is essential to ensure homeostasis of the organism and coordinate fundamental behaviors, such as feeding, mating and reproduction, and acts as integrator of external and internal sensory cues [33]. Consequently, connections between sensory and endocrine systems are found across species [34–37]. We show that hugin neurons receive chemosensory input in the *Drosophila* analog of the brain stem, thereby linking chemosensory and neuroendocrine systems. Overall, these findings reveal evolutionary constancy among the neural circuits of hugin and NMU.

## Results

### Input and output compartments of hugin neurons

The hugin gene is expressed in only 20 neurons in the CNS. This population comprises interneurons, which are confined within the CNS, as well as efferent neurons, which leave the CNS. The interneuron type can be subdivided into those projecting to the protocerebrum (hugin-PC, 8 neurons) or the ventral nerve cord (hugin-VNC, 4 neurons).

The efferent type can be subdivided into those projecting to the ring gland (hugin-RG, 4 neurons) or the pharynx (hugin-PH, 4 neurons) (Fig. 2A) [20]. Based on these morphological features, we first reconstructed all hugin neurons in an ssTEM volume covering an entire larval CNS and the major neuroendocrine organ, the ring gland (Fig. 2B; see materials and methods for details). We then localized synaptic sites, which could be readily identified as optically dense structures [38]. Comparing neurons of the same class, we found the number as well as the distribution of pre- and postsynaptic sites to be very similar among hugin neurons of the same class (Fig. 2C–E, Video 1). Presynaptic sites are generally defined as having small clear core vesicles (SCVs) containing classic small molecule transmitter for fast synaptic transmission close to the active zone [38]. Efferent hugin neurons (hugin-RG and hugin-PH) showed essentially no presynaptic sites (<1 average/neuron) within the CNS and we did not observe any SCVs. For hugin-RG neurons, peripheral presynaptic sites were evident at their projection target, the ring gland. These presynaptic sites did contain close-by DCVs but no SCVs and had in many cases no corresponding postsynaptic sites in adjacent neurons. Instead they bordered haemal space indicating neuroendocrine release (Fig. 2 – figure supplement 1 A). The configuration of hugin-PH terminals is unknown as their peripheral target was outside of the ssTEM volume. For the interneuron classes (hugin-PC and hugin-VNC), we found SCVs at larger presynaptic sites, indicating that they employ classic neurotransmitter in addition to the hugin peptide (Fig. 2 – figure supplement 1 B,C). Hugin-PC and hugin-VNC neurons’ projections represent mixed synaptic input-output compartments as they both showed pre- as well as postsynaptic sites along their neurites (Fig. 2D, E).

**Figure 2.**
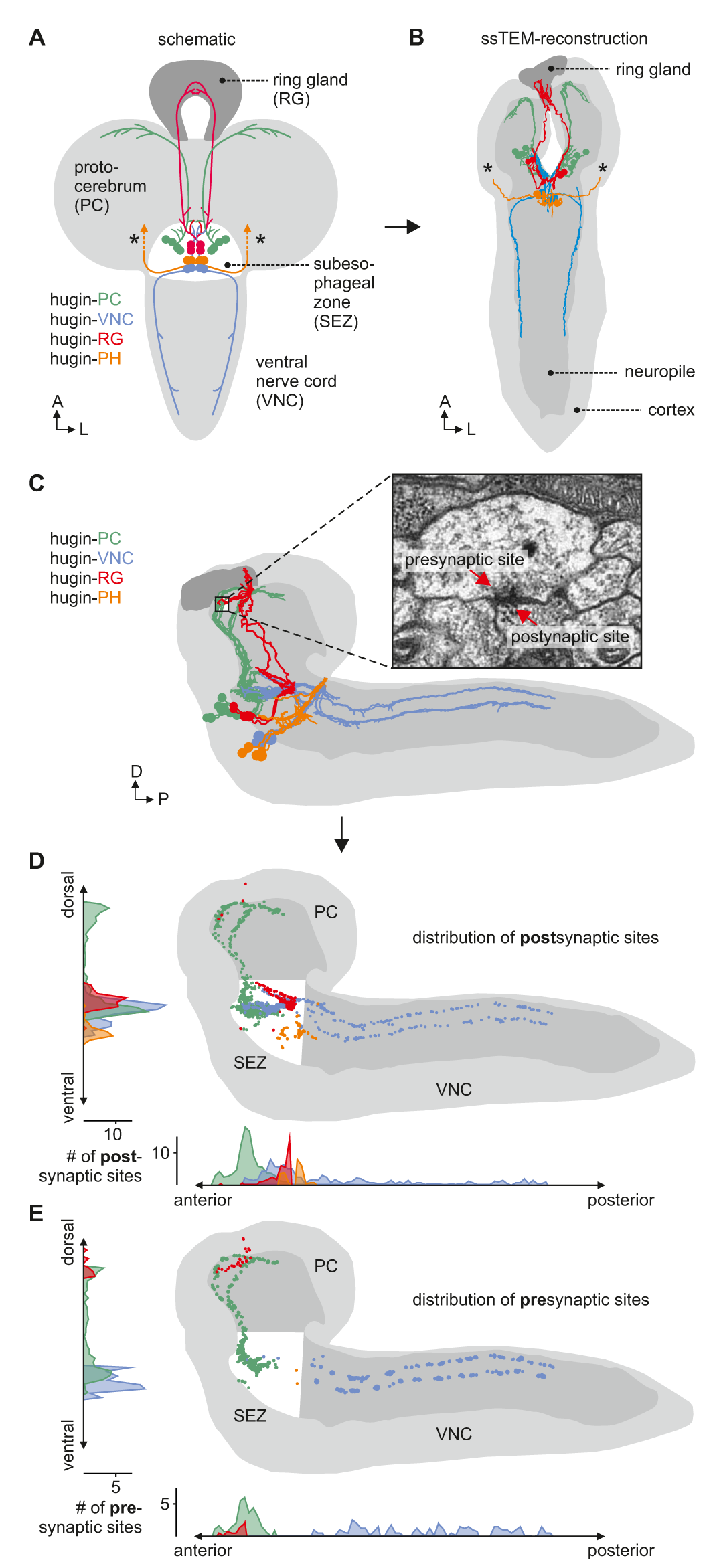
Morphology of hugin-producing neurons and their input and output compartments. **A**, Four morphologically distinct classes of hugin neurons: hugin-PC (protocerebrum), hugin-VNC (ventral nerve cord), hugin-RG (ring gland) and hugin-PH (pharynx, asterisks mark nerve exit sites). **B**, Reconstruction of all hugin neurons based on an entire larval brain. **C**-**E**, Spatial distribution of pre-and postsynaptic sites for all hugin classes. Each dot in D and E represents a single synaptic site. Graphs show distribution along dorsal-ventral and anteriorposterior axis of the CNS. Hugin interneurons (hugin-PC and hugin-VNC) show mixed input and output compartments, whereas efferent hugin neurons (hugin-RG and hugin-PH) show almost exclusively postsynaptic sites within the CNS. Note that presynaptic sites of hugin-RG neurons (E) are located in the ring gland. See also video 1.

All hugin neurons receive inputs within the subesophageal zone [SEZ, previously called subesophageal ganglion (SOG)], a chemosensory center that also houses the basic neuronal circuits generating feeding behavior [39]. However, only the hugin-PC neurons showed considerable numbers of synaptic outputs in the SEZ, consistent with their previously reported effects on feeding [19,40] (Fig. 2E).

### Acetylcholine is a co-transmitter in hugin neurons

The existence of presynaptic sites containing SCVs in addition to large DCVs led to the assumption that hugin-PC and hugin-VNC (possibly also hugin-PH neurons) employ small molecule neurotransmitters in addition to the hugin neuropeptide. To address this, we checked for one of the most abundantly expressed neurotransmitter in the *Drosophila* nervous system: acetylcholine (ACh) [41,42]. In the past, immunohistochemical and promoter expression analyses of choline acetyltransferase (ChAT), the biosynthetic enzyme for ACh, were successfully used to demonstrate cholinergic transmission [43–45]. We used both, anti-ChAT antibody as well as a ChAT promoter GAL4 driving expression of a fluorescent reporter and investigated co-localization with hugin neurons. In EM data hugin neurons had comparatively few SCVs suggesting only low amounts of small transmitters. In addition, ChAT is preferentially localized in the neuropil and less so in the somas [46]. Consistent with this, we found that ChAT immunoreactivity in hugin cell bodies was relatively low and varied strongly between samples. Therefore we quantified the anti-ChAT signal to show that while ChAT levels were in some cases indiscernible from the background, overall highest levels of ChAT were found in hugin-PC and hugin-VNC/PH neurons (Fig. 3A). Note that while hugin-PC and hugin-RG neurons were easily identifiable based on position and morphology, hugin-PH and hugin-VNC neurons usually clustered too tightly to be unambiguously discriminated and were thus treated as a single mixed group. Similar to the immunohistochemical analysis, the ChAT promoter (ChAT-GAL4) drove expression in all hugin-PC neurons plus subset of hugin-VNC/PH neurons (Fig. 3B). Hugin-RG showed weak ChAT signal with either method, consistent with these neurons lacking SCVs in the EM data.

**Figure 3.**
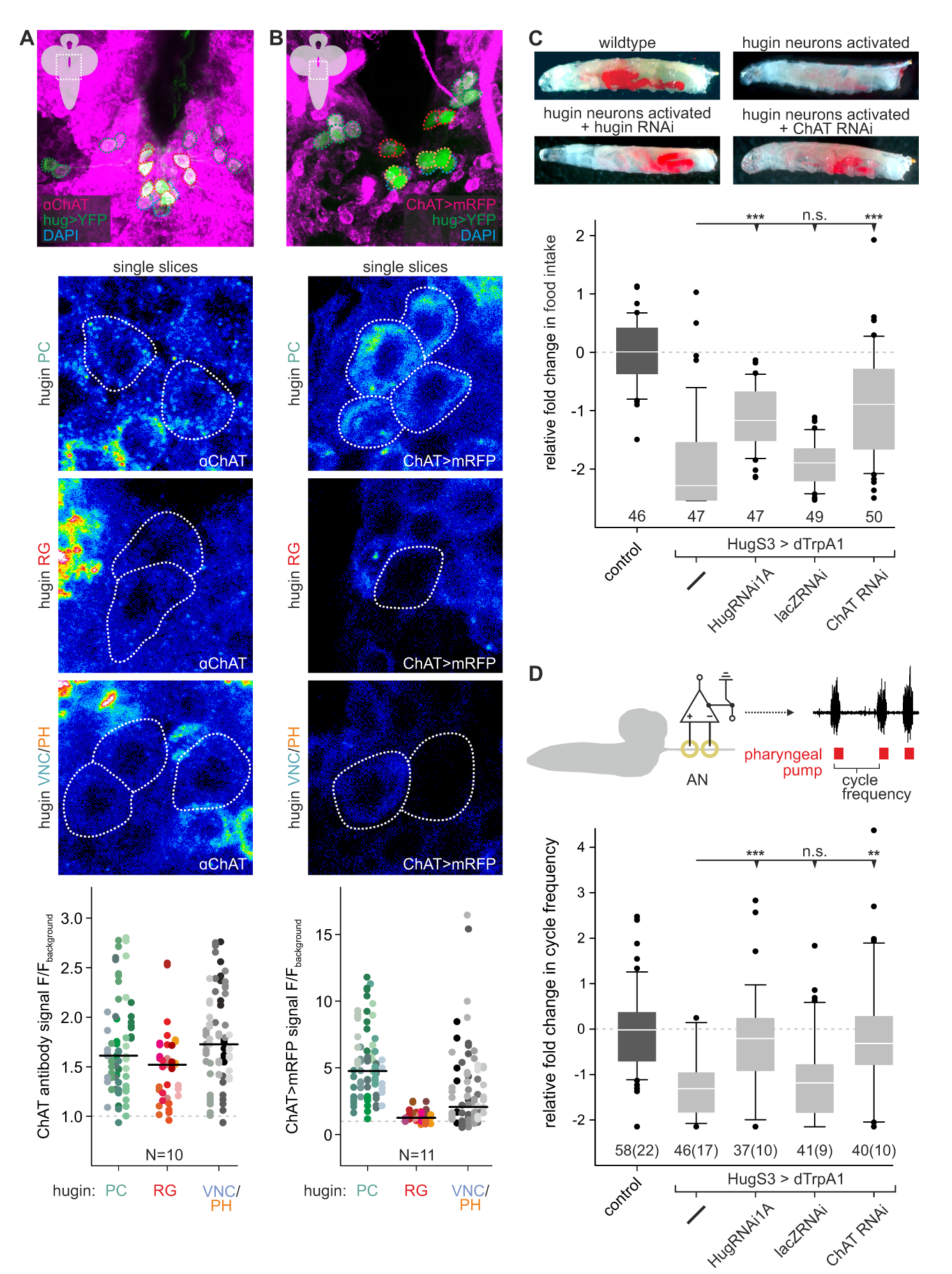
Acetylcholine (ACh) is a neurotransmitter of hugin neurons. **A,B**, Co-localization of the biosynthetic enzyme for ACh, choline acetyltransferase (ChAT), in hugin neurons using a ChAT antibody (A) or a ChAT promoter-GAL4 driving a fluorescent reporter (B). Shown are exemplary scans and quantification of ChAT co-localization in the different hugin classes using the two approaches. Each data point in dot plots represents a single hugin neuron. Horizontal line marks median. ChAT immunoreactivity was variable but strongest signals were found in hugin-PC and hugin-VNC/PH neurons. Similarly, ChAT-GAL4 consistently drove expression in hugin-PC and subsets of hugin-VNC/PH. Note that while hugin-PC and hugin-RG neurons are easily identifiable, hugin-PH and hugin-VNC neurons were usually too close to be unambiguously discriminated and were thus treated as a single mixed group. **C**,**D**, Contribution of ACh to hugin effect. Food intake (C) and extracellular recordings (D) of the antennal nerve (AN). AN recordings were analyzed in respect to the cycle frequency of the fictive motor activity of the pharyngeal pump. In both assays, hugin neurons were artificially activated using the thermosensitive cation channel dTrpA1. Decrease in food intake and pharyngeal pump activity due to activation of hugin neurons was rescued by RNAi-induced knockdown of either the hugin neuropeptide or ChAT. Data in C and D taken from Schoofs et al. (2014 [19]; Fig. 5B, C) and extended by ChAT RNAi experiments. Numbers below box plots give N (C, # larvae; D, # trials (# larvae). Mann-Whitney Rank Sum Test (*** = p<0.001;**=p<0.01).

These findings suggested that ACh may be a co-transmitter in hugin neurons. We previously demonstrated that RNAi-induced knockdown of the hugin neuropeptide rescues the phenotype of feeding suppression caused by induced activation of hugin neurons in behavioral and electrophysiological experiments [19]. Here, we extended this initial data by a knockdown of ChAT using an established UAS-ChAT-RNAi line [47]. We found that knockdown of ChAT in hugin neurons rescued the decrease in food intake to a similar degree as the knockdown of the hugin neuropeptide itself (Fig. 3C). This rescue by knockdown of either hugin or ChAT was confirmed using extracellular recordings of the antennal nerve (AN) in isolated CNS for precise monitoring of motor pattern of the pharyngeal pump (Fig. 3D) [48]. Activation of hugin neurons leads to a decrease in cycle frequency of pharyngeal pump motor activity. This too was rescued by either hugin neuropeptide or ChAT knockdown. Taken together, this data clearly demonstrates that ACh plays a functional role in hugin neurons. Moreover, it suggests that hugin neuropeptide and ACh have to be employed together in order to regulate feeding behavior.

### Hugin classes form distinct units that share synaptic partners

Reconstruction of hugin neurons and localization of synaptic sites revealed that neurons of the two interneuron classes, hugin-PC and hugin-VNC, were reciprocally connected to ipsilateral neurons of the same class (Fig. 4, Fig. 2 – figure supplement 1 E,F). These axo-axonic synaptic connections made up a significant fraction of each neuron’s synaptic connections, implying that their activity might be coordinately regulated.

**Figure 4.**
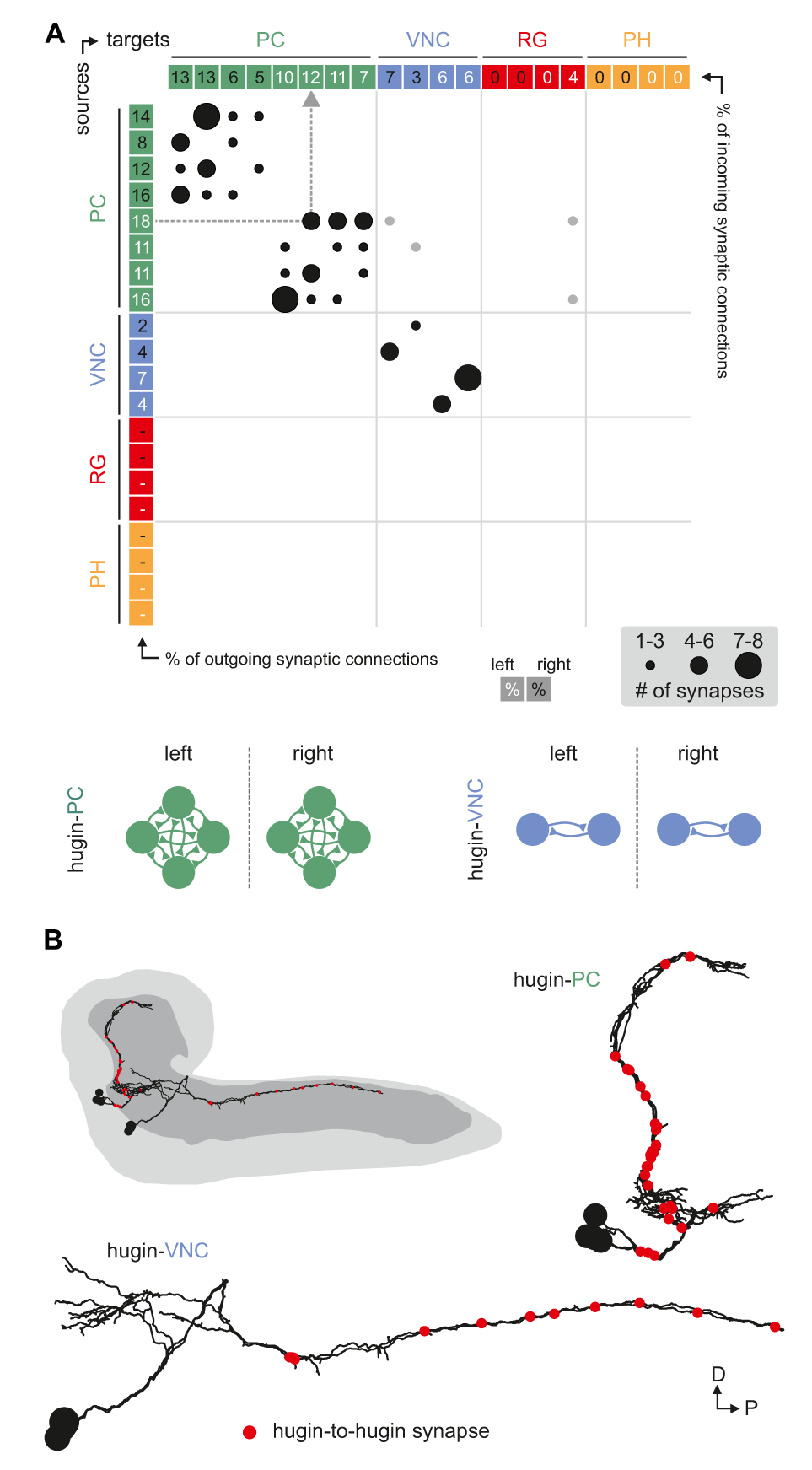
Hugin neurons synapse axo-axonically reciprocally within-class but not across-class. **A**, Connectivity matrix of hugin to hugin connections. Each row indicates number of synaptic connections of given hugin neuron to other hugin neurons. Connections that could not be recapitulated for both hemisegments are grayed out. Numbers in colored boxes give % of incoming (x-axis) and outgoing (y-axis) synaptic connections of the respective hugin neuron. Hugin to hugin contacts are made between hugin interneurons of the same class, not between classes (see schematic). Note that efferent hugin neurons, hugin-RG and hugin-PH, do not have presynaptic sites. **B**, Distribution of hugin-hugin synapses. Synapses connecting hugin-PC to hugin-PC and hugin-VNC to hugin-VNC neurons are axi-axonic. Only neurons of one hemisegment are shown.

We therefore further explored the different classes within the population of hugin-producing neurons, asking whether hugin classes establish functional units or whether they are independently wired. To this end, we reconstructed 177 pre- and postsynaptic partners of hugin neurons (Fig. 5A, see materials and methods for details). First, we found that neurons of the same hugin class were connected to the same pre- and postsynaptic partners. Furthermore, most synaptic partners were connected exclusively to neurons of a single hugin class (Fig. 5B). Second, pre- and postsynaptic partners of each hugin class resided in different parts of the CNS (Fig. 5C; Video 2). For hugin-RG and hugin-PH the vast majority of synapses were made with interneurons, 93±4% and 97±3%, respectively. This percentage was lower for hugin-PC (66±6%) and hugin-VNC (81±2%). To our knowledge, none of these reconstructed interneuron partners have been previously described, making it difficult to deduce functional implications at this point. Non-interneuron partners will be described in the following sections. In summary, these findings show that neurons of each hugin class form complex microcircuits that are largely separate from one another.

**Figure 5.**
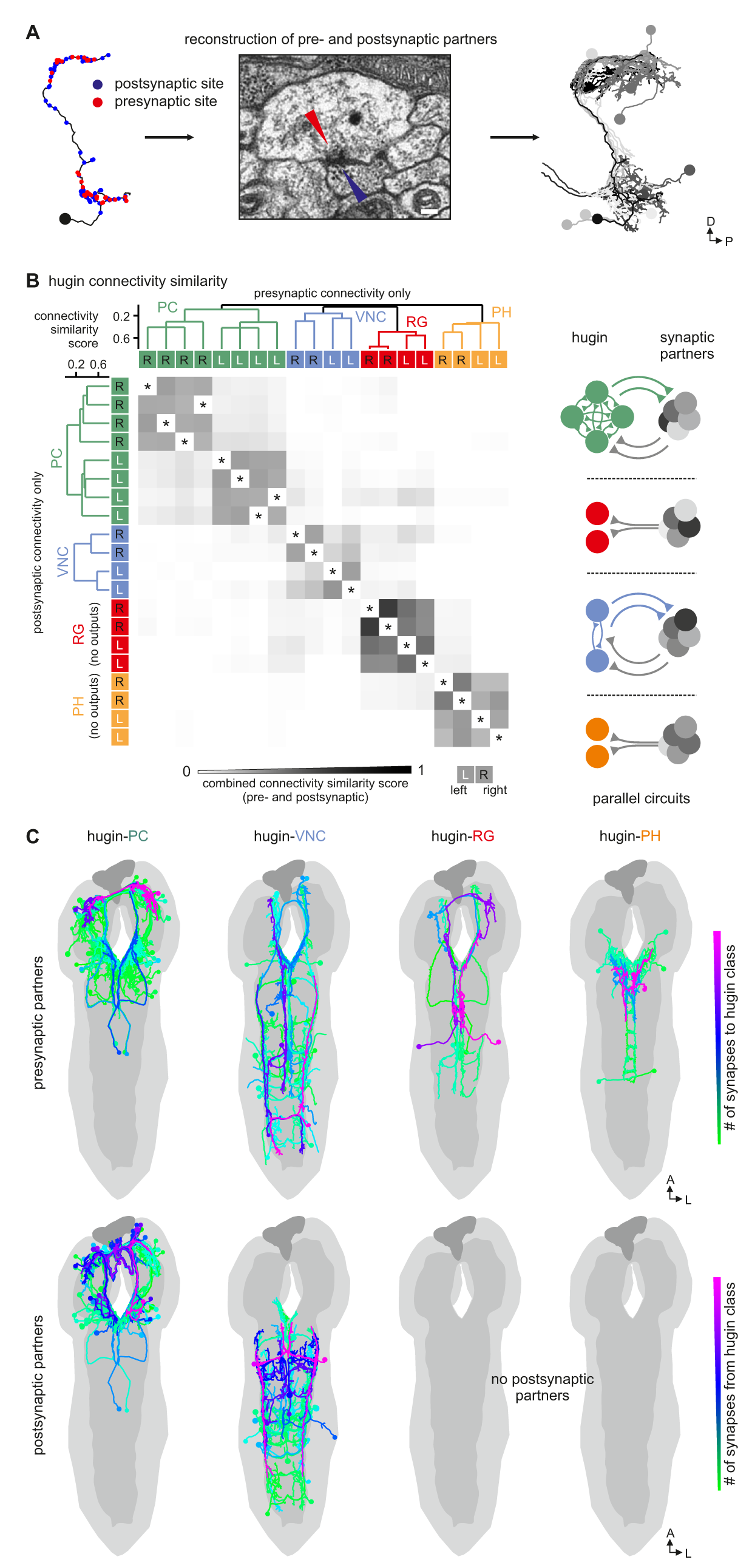
Each hugin class is part of a distinct microcircuit, weakly or not at all connected to those of the other classes. **A**, Exemplary pre- and postsynaptic partners of a single hugin neuron. **B**, Overlap of synaptic partners of individual hugin neurons as measured by connectivity similarity score. High similarity score indicates a large fraction of shared synaptic partners connected by similar numbers of synapses. Neurons are ordered by dendrogram of similarity score of pre- (x-axis) and postsynaptic (y-axis) partners. Matrix shows combined pre- and postsynaptic similarity score. Self-self comparisons were omitted (asterisks). Hugin classes connect to unique sets of pre- and postsynaptic partners. Neurons of each hugin class have the same synaptic partners and there is little to no overlap with other classes (see schematic). **C**, All pre- and postsynaptic partners by hugin class. Neurons are color-coded based on total number of synapses to given hugin class [minimum=1; maximum (pre-/postsynaptic):hugin-PC=53/16, hugin-VNC=21/18, hugin-RG=39/none,hugin-PH=23/none]. Hugin-RG and hugin-PH neurons do not have postsynaptic partners within the CNS. See also video 2.

### Hugin neurons receive diverse chemosensory synaptic input

Hugin neurons have a significant number of their incoming synapses (63 ± 22%) within the SEZ. This region of the CNS is analogous to the brainstem and is a first order chemosensory center that receives input from various sensory organs [24]. In addition, subsets of hugin neurons were recently shown to be responsive to gustatory stimuli [40]. We therefore searched for sensory inputs to hugin neurons and found a total of 68 afferent neurons that made synaptic contacts onto hugin neurons (Fig. 6A). Two major groups emerged: a larger, morphologically heterogeneous group consisting of afferent neurons projecting through one of the pharyngeal nerves (the antennal nerve) and, unexpectedly, a second, more homogeneous group entering the CNS through abdominal (but not thoracic) nerves. We observed that the reconstructed afferent presynaptic partners of hugin neurons covered different parts of the SEZ. Thus, we sought to cluster these afferent neurons by computing the similarity in spatial distribution of their synaptic sites, termed synapse similarity score.

**Figure 6.**
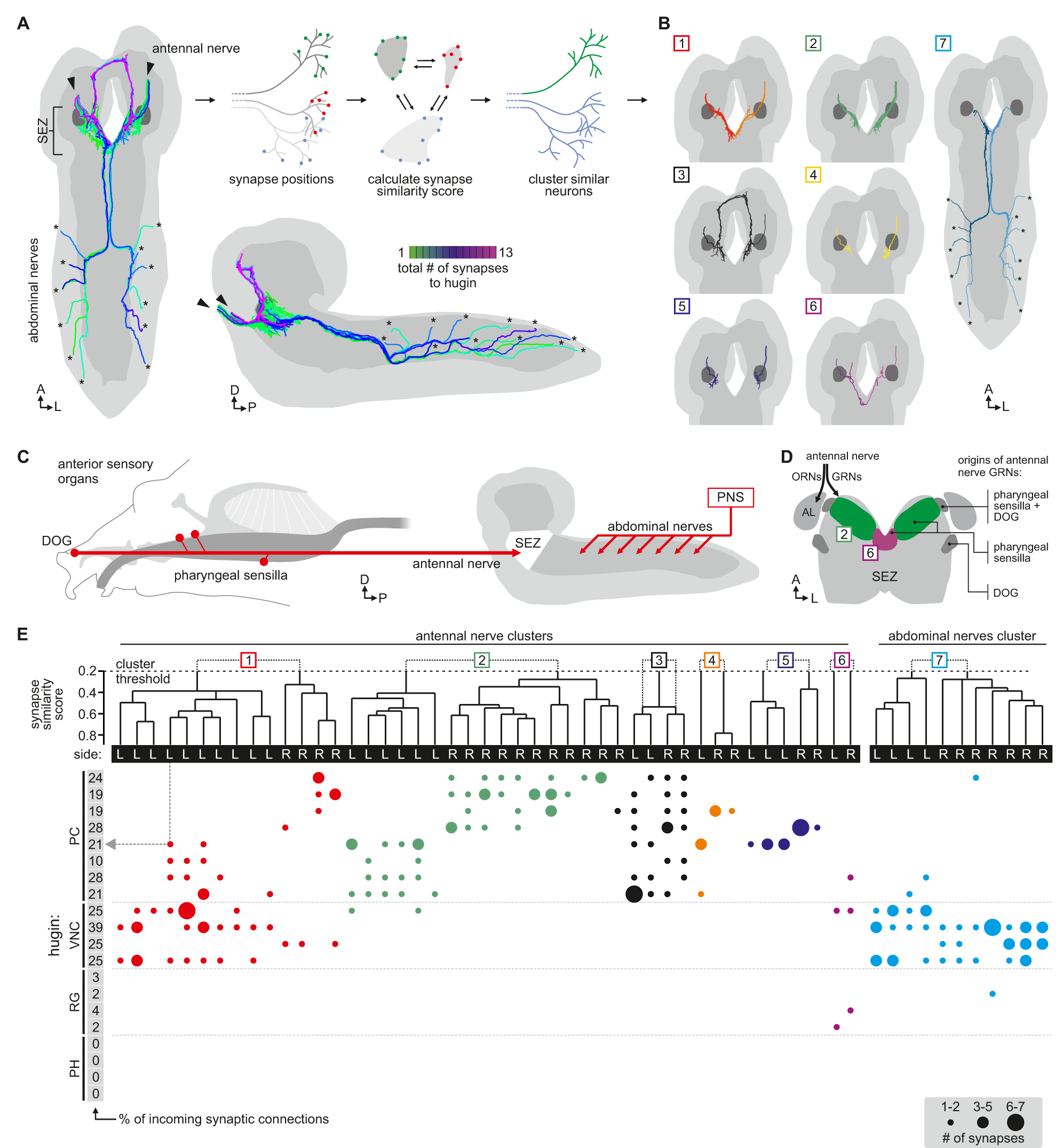
Each class of hugin neurons receives inputs from distinct subsets of sensory neurons. **A**, Sensory inputs to hugin neurons enter the CNS via the antennal nerve (arrowheads) and abdominal nerves (asterisks). Neurons are color-coded based on total number of synapses to hugin neurons. **B**, Morphology of sensory neurons clustered based on a synapse similarity score computed from the spatial overlap of synaptic sites between two neurons. See also video 3. **C**, Potential origins of sensory inputs onto hugin neurons. The antennal nerve collects sensory axons from the dorsal organ ganglion (DOG) and pharyngeal sensilla. Abdominal nerves carry afferents from the abdominal segments of the peripheral nervous system (PNS). **D**, Target areas of antennal nerve chemosensory organs in the subesophageal zone (SEZ). Olfactory receptor neurons (ORNs) terminate in the antennal lobes (AL). Gustatory receptor neurons (GRNs) from different sensory organs cover distinct parts of the SEZ (based on Colomb et al.,2007). **E**, Connectivity matrix of sensory neurons onto hugin. Sensory neurons are ordered by dendrogram of synapse similarity score and rearranged to pair corresponding cluster of left (L) and right (R) hemisegment. Each row of the matrix shows the number of synaptic connections onto a single hugin neuron. Numbers in gray boxes along y-axis give percentage of synaptic input onto each hugin neuron represented as one neuron per row. Only sensory neurons that have at least a single more-than-2-synapse connection to hugin neurons are shown. See text for further details.

Clustering based on synapse similarity score resulted in 7 different groups, each of them covering distinct parts of the SEZ (Fig. 6B; Video 3; see material and methods for details). To address the issue of the origin of identified sensory inputs, we compared our data with previous descriptions of larval sensory neurons. It is well established that abdominal nerves innervate internal and external sensory organs of the peripheral nervous system. This includes proprioceptive (chordotonal), tactile, nociceptive (multi dendritic neurons) and a range of sensory neurons whose function is yet unknown [49–51]. To our knowledge no abdominal sensory neurons with projections into the SEZ such as the one observed presynaptic to hugin have been described. However, the majority of afferent neurons synapsing onto hugin neurons stems from the antennal nerve. This pharyngeal nerve carries the axons of gustatory receptor neurons (GRNs) from internal pharyngeal sensilla as well as those of olfactory receptor neurons (ORNs) and other GRNs from the external sensory organs (Fig. 6C,D) [52,53]. ORNs can be unambiguously identified as they target specific glomeruli of the antennal lobe [53] but no such sensory neurons were found to directly input onto hugin neurons (Fig. 6 – figure supplement 1).

The GRNs likewise target restricted regions of the SEZ neuropil but this is not as well characterized as the antennal lobes [52,54]. The antennal nerve neurons that contact the hugin cells show the morphology of this large, heterogeneous population of GRNs [52,55]. We thus compared our clustered groups with previously defined light microscopy-based gustatory compartments of the SEZ [52]. Groups 2 and 6, which cover the anterior-medial SEZ, likely correspond to two areas described as the target of GRNs from internal pharyngeal sensilla only (Fig. 6D). The remaining groups were either not previously described or difficult to unambiguously align with known areas. Our division into groups is also reflected at the level of their connectivity to hugin neurons: sensory neurons of group 1 have synaptic connections to both hugin-PC and hugin-VNC neurons. Groups 2-5, encompassing the previously described pharyngeal sensilla, are almost exclusively connected to hugin-PC neurons. Group 6 sensory neurons make few synapses onto hugin-RG neurons. Group 7, encompassing the abdominal afferent neurons, is primarily presynaptic to hugin-VNC (Fig. 6E).

The efferent type hugin neurons, hugin-PH and hugin-RG, show little to no sensory input. In contrast, the interneuron type hugin neurons, hugin-PC and hugin-VNC, receive a significant fraction of their individual incoming synaptic connections (up to 39%) from sensory neurons. Summarizing, we found two out of four types of hugin neurons to receive synaptic input from a large heterogeneous but separable population of sensory neurons, many of which are GRNs from external and internal sensory organs. Hugin-PC neurons were recently shown to be activated by bitter gustatory stimuli but not salt, fructose or yeast [40]. Our data strongly indicates that this activation is based on monosynaptic connections to GRNs. Moreover, the heterogeneity among the population of sensory neurons suggests that hugin-PC neurons do not merely function as simple relay station but rather fulfill an integrative function, for example between multiple yet-to-be-identified modalities or various external and internal sensory organs.

### Dual synaptic and peptide-receptor connection to the neuroendocrine system

NMU has been well studied in the context of its effect on the hypothalamo-pituitary axis. We therefore looked for similar motifs among the downstream targets of hugin neurons. The cluster of hugin-PC neurons projects their neurites from the SEZ to the protocerebrum, terminating around the pars intercerebralis. Median neurosecretory cells (mNSCs) in this area constitute the major neuroendocrine center in the CNS, homologous to the mammalian hypothalamus, and target the neuroendocrine organ of *Drosophila*, the ring gland [25].

Three different types of mNSCs produce distinct neuropeptides in a non-overlapping manner: 3 mNSCs produce diuretic hormone 44 (DH44), 2 mNSCs produce Dromyosuppressin (DMS) and 7 mNSCs produce *Drosophila* insulinlike peptides (Dilps, thus called insulin-producing cells [IPCs]) (Fig. 7A) [56]. We found that hugin-PC neurons make extensive synaptic connections onto most but not all of the mNSCs (Fig. 7B; Fig. 2 – figure supplement 1 G,H). mNSCs of the pars intercerebralis derive from the same neuroectodermal placodes and develop through symmetric cell division [57]. Among the mNSCs, IPCs have been best studied: they have ipsilateral descending arborizations into the SEZ and project contralaterally into the ring gland [2]. In contrast, morphology of DH44- or DMS-producing mNSCs has been described in less detail. Our reconstruction showed that all reconstructed mNSCs have the exact same features, rendering them morphologically indistinguishable (Fig. 7C). To assign identities to the reconstructed mNSCs, we hypothesized that similar to hugin neurons, the three types of mNSCs would differ in their choice of synaptic partners. We therefore reconstructed all presynaptic partners and calculated the connectivity similarity score between the mNSCs. Clustering with this similarity in connectivity resulted in three groups comprising 3, 2 and 7 neurons, coinciding with the number of neurons of the known types of mNSCs. We thus suggest that the group of 3 represents DH44-producing cells, the group of 2 represents DMS-producing cells and the group of 7 represents the IPCs (Fig. 7D).

**Figure 7.**
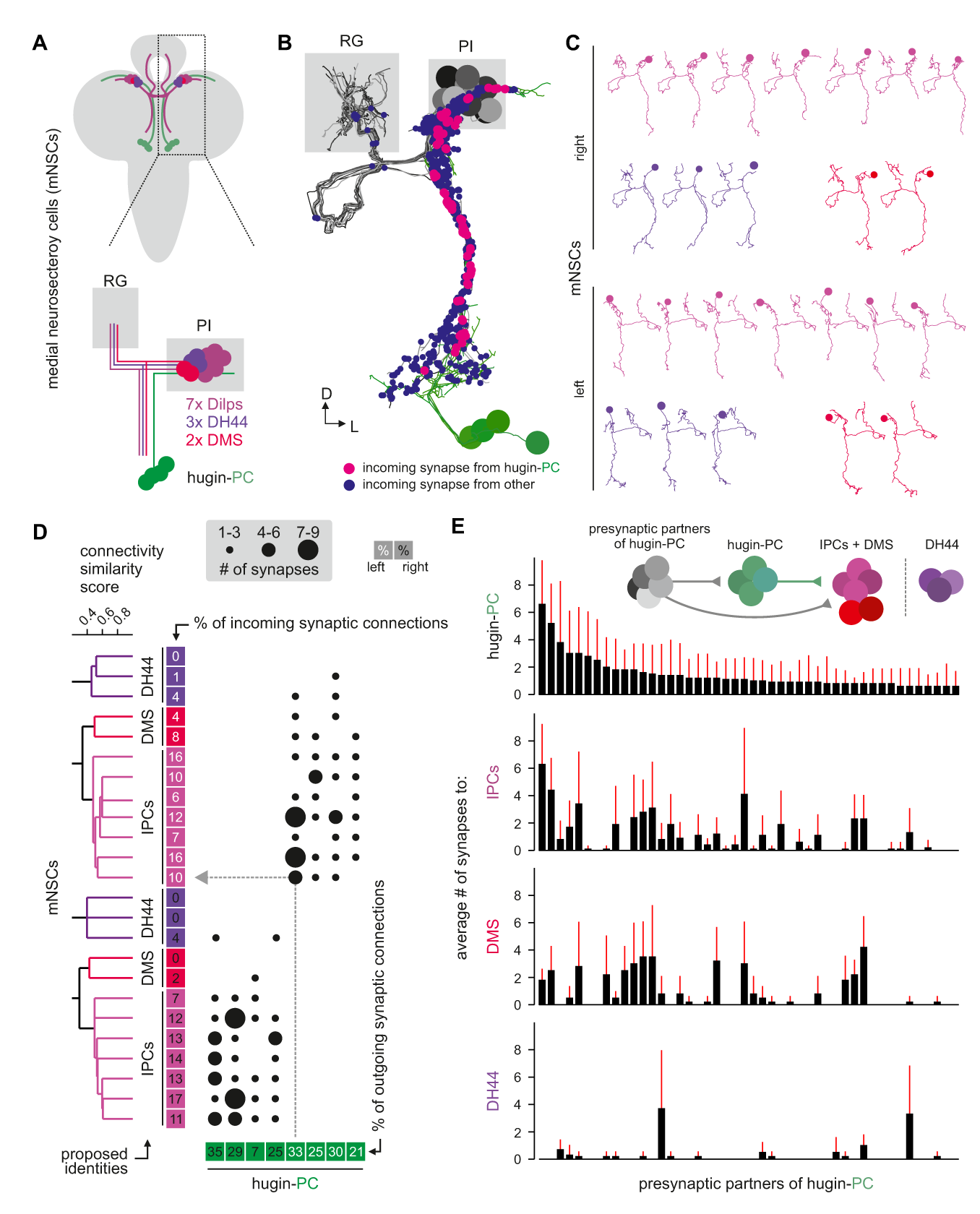
Hugin-PC neurons are presynaptic to all insulin-producing neurosecretory neurons. **A**, Schematic of median neurosecretory cells (mNSCs) located in the pars intercerebralis (PI). mNSCs produce *Drosophila* insulin-like peptides (Dilps), diuretic hormone 44 (DH44) and Dromyosuppressin (DMS). **B**, EM-reconstruction of all mNSCs. Hugin-PC neurons make axo-axonic synapses onto mNSCs. **C**, All mNSCs are sibling neurons derived from the same neuroectodermal placode via symmetric cell division. Ipsilateral siblings present similar arborizations, making morphological identification impossible. Instead mNSCs were categorized by connectivity (see D). **D**, Connectivity matrix of hugin-PC to mNSCs. mNSCs are ordered by dendrogram of connectivity similarity of all presynaptic partners. Proposed identity is based on connectivity similarity clustering into groups of 3 DH44-, 2 DMS- and 7 Dilps-producing cells (see text for details). In the matrix, each row indicates number of synapses from a single hugin-PC neuron onto a mNSCs. Numbers in boxes give % of outgoing/incoming synaptic connections represented in each column or row, respectively. **E**, Connectivity between presynaptic partners of hugin-PC neurons and mNSCs. Each column across all four graphs represents a presynaptic partner of hugin-PC. Graphs show average number of synapses to hugin-PC, DH44-, DMS- and Dilps-producing neurons of given presynaptic partner. Whiskers represent standard deviation. hugin-PC neurons share inputs with Dilps- and DMS-producing neurons but not with DH44-producing neurons.

On this basis, hugin-PC neurons make extensive synaptic contacts to the IPCs but less so to DMS- and DH44-producing mNSCs. In accordance with hugin-PC neurons using ACh as neurotransmitter, IPCs were previously shown to express a muscarinic ACh receptor [58]. Overall, synapses between hugin-PC neurons onto mNSCs constitute a large fraction of their respective synaptic connections (hugin-PC: up to 35%; mNSCs: up to 17%). In support of a tight interconnection between hugin neurons and these neuroendocrine neurons, we found that most of hugin-PC neurons’ presynaptic partners are also presynaptic to mNSCs (Fig. 7E). These findings demonstrate that the neuroendocrine system is a major target of hugin neurons.

Unlike the small molecule messengers used for fast synaptic transmission, neuropeptides - such as hugin - are thought to be released independent of synaptic membrane specializations and are able to diffuse a considerable distance before binding their respective receptors. However, it has been proposed that neuropeptides released from most neurons act locally on cells that are either synaptically connected or immediately adjacent (van den Pol 2012). We therefore asked whether the synaptic connections between hugin-PC neurons and mNSCs would have a matching peptide-receptor connection. The hugin gene encodes a prepropeptide that is post-translationally processed to produce an eight-amino-acid neuropeptide, termed pyrokinin-2 (hug-PK2) or hugin neuropeptide (Meng et al. 2002). This hugin neuropeptide has been shown to activate the *Drosophila* G-Protein coupled receptor (GPCR) CG8784/PK2-R1 in mammalian cell systems, but the identities of the target neurons expressing the receptor remain unknown (Rosenkilde et al. 2003). To address this, we used two independent methods to generate transgenic fly lines, CG8784-GAL4::p65 and CG8784-6kb-GAL4, driving expression under control of putative CG8784 regulatory sequences (Fig. 8A; Fig. 8 – figure supplement 1). Both CG8784-GAL4 lines drive expression of a GFP reporter in a prominent cluster of cells in the pars intercerebralis. Double stainings show that this expression co-localizes with the peptides produced by the three types of mNSCs: Dilp2, DH44 and DMS (Fig. 8B–D; Fig. 8 – figure supplement 1). To support the receptor expression data, we performed calcium imaging of the mNSCs upon treatment with hug-PK2 (Fig. 8 – figure supplement 2). Indeed, calcium activity of the mNSCs increased significantly after treatment with concentrations of 1 μM hug-PK2 or higher. These findings indicate that hugin-PC neurons employ both classical synaptic transmission and peptidergic signaling to target neurons of the neuroendocrine center.

**Figure 8.**
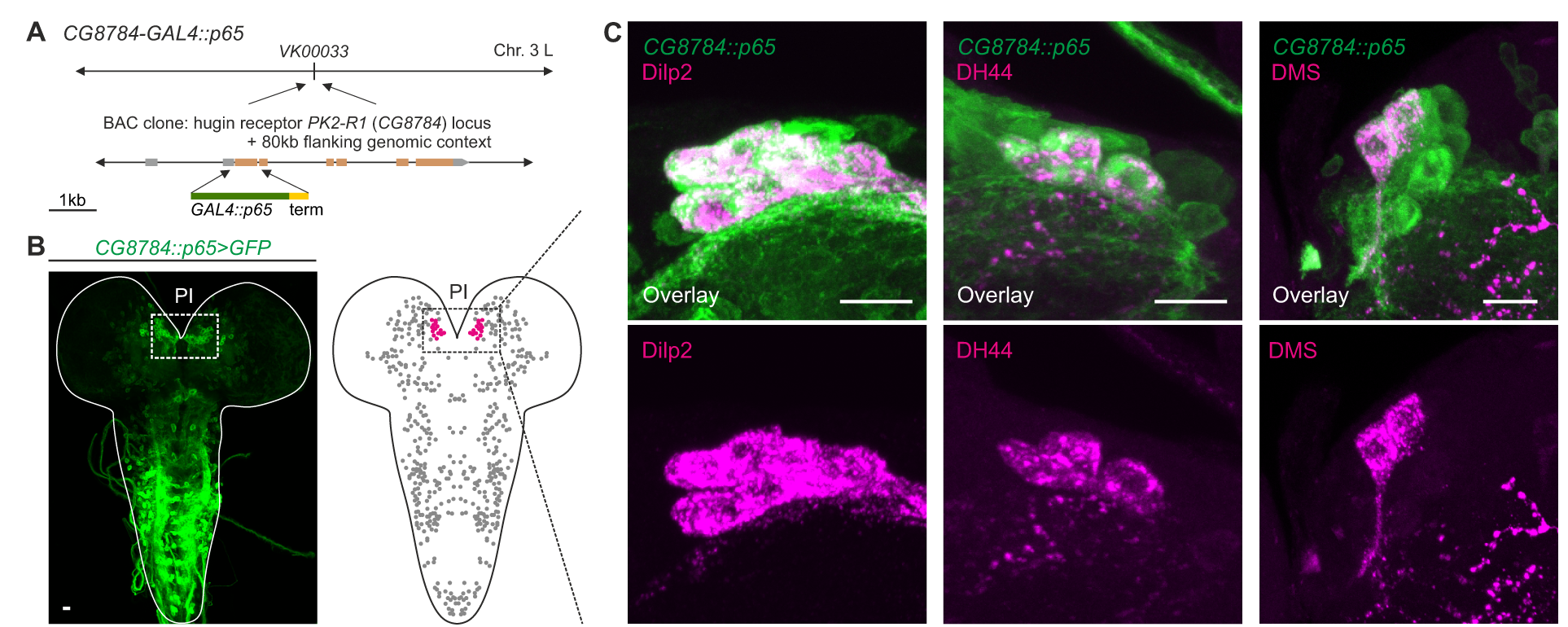
GPCR-mediated neuromodulatory transmission is used in addition to synaptic connections. **A**, Promoter-based hugin G-protein coupled receptor (GPCR) PK2-R1 driver line CG8784-GAL4::p65 was generated by replacing the first coding exon of the CG8784 loci with GAL4 in a BAC clone containing 80kb flanking genomic context and integrating the final BAC into attP site VK00033. **B**, CG8784-GAL4::p65 drives expression in cells of the pars intercerebralis (PI). **C**, Co-staining with *Drosophila* insulin-like peptide 2 (Dilp2), diuretic hormone 44 (DH44) and Dromyosuppressin (DMS). These peptides are produced by median neurosecretory cells (mNSCs) in a non-overlapping manner. CG8784-GAL4::p65 drives expression in all mNSCs of the PI. Scale bars in A and B represent 10 μm.

Neuropeptides are produced in the soma and packaged into dense core vesicles (DCVs) before being transported to their release sites [59]. We found 98% of the synapses between hugin-PC and mNSCs to have DCVs within 1000 nm radius to the presynaptic sites (average # of DCVs/synapse: 15.5), opening up the possibility of co-transmission of peptide and classical neurotransmitter [60] (Fig. 9A). Further exploring the spatial relationship between DCVs and synapses, we observed that for both interneuron type hugin classes (hugin-PC and hugin-VNC) DCVs localized close to presynaptic sites. This was often the case at local swellings along the main neurites which featured multiple pre- and postsynaptic sites, as well as close-by DCVs (Fig. 9B,C). It is conceivable that such complex local synaptic circuitry might enable local peptide release. Next, we measured the distance to the closest presynaptic site for each DCV. The majority of DCVs in hugin-PC and hugin-VNC neurons was localized within approximately 2000 nm from the next presynaptic site (Fig. 9D). However, most DCVs were probably too distant from presynaptic sites to be synaptically released, suggesting para- and non-synaptic release [61,62] (Fig. 2 – figure supplement 1 D).

**Figure 9.**
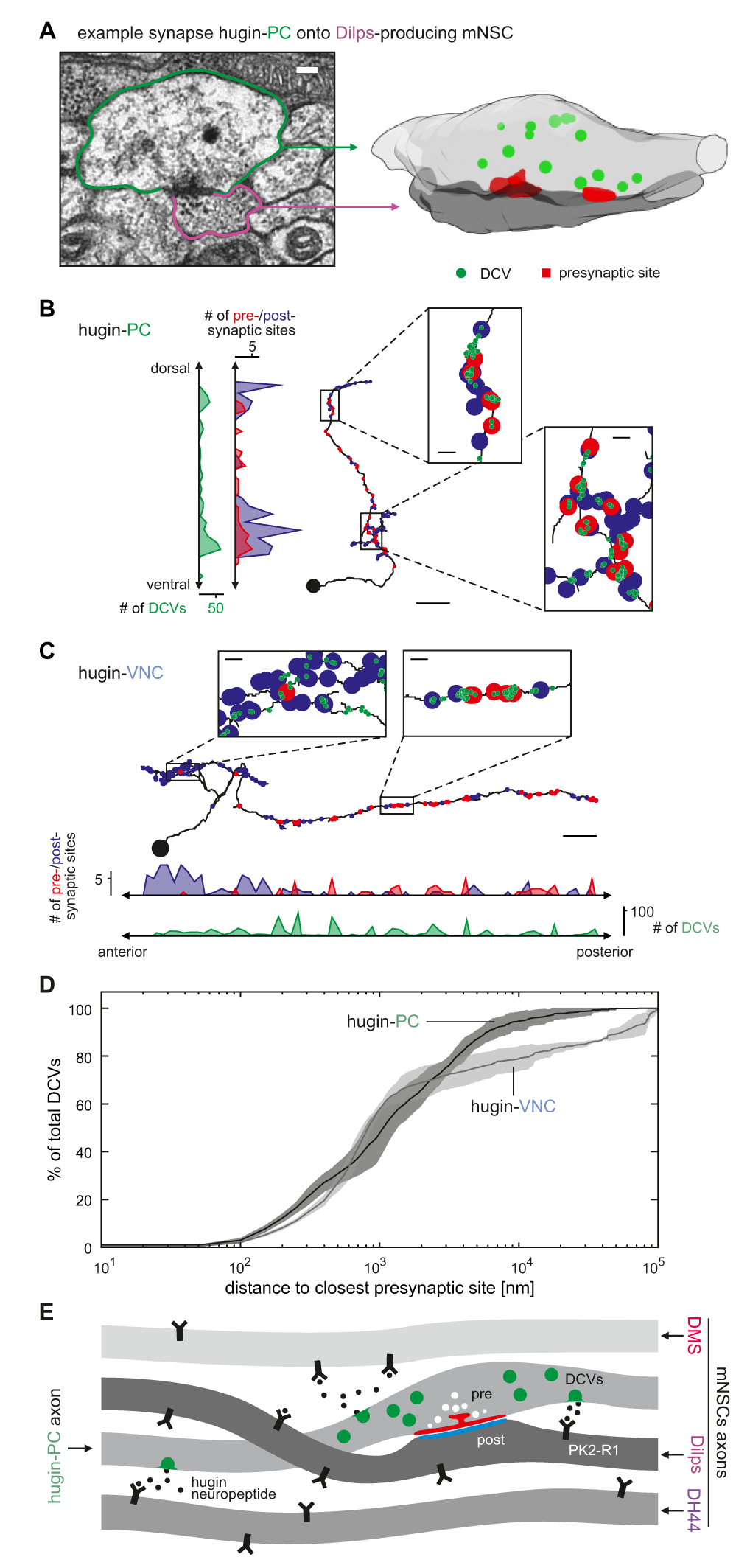
Dense core vesicles localize close to but not directly at presynaptic sites. **A**, Volume reconstruction of example synapse between hugin-PC neuron and median neurosecretory cells (mNSCs) producing *Drosophila* insulinlike peptides (Dilps) shows dense core vesicles (DCV) in the vicinity of synaptic densities. Scale bar represents 1OO pm. **B**,**C**, Distribution of pre- and postsynaptic sites and DCVs for a single hugin-PC (B) and hugin-VNC (C) neuron. DCVs localize close to presynaptic sites. Scale bars represent 1O pm (overview) and 1 μm (inlets). **D**, For each DCV the distances to the closest presynaptic was calculated. Graph shows percentage of DCVs within given distance to closest presynaptic site. Vesicles in the soma and the proximal part of the main neurite were excluded. Only a small fraction of DCVs are in very close proximity to presynaptic sites, indicating para- and non-synaptic rather than synaptic release. Note that hugin-RG and hugin-PH neurons were excluded due to lack of presynaptic sites within the CNS. Dashed vertical lines mark 50% fraction. Envelopes represent standard deviation. **E**, Summarizing schematic and model. Hugin-PC neurons make classical chemical synapses almost exclusively onto Dilps-producing mNSCs. Additionally, all mNSCs express hugin receptor PK2-R1 (CG8784) and are often in close vicinity to hugin neurites, allowing para- or non-synaptically released hugin neuropeptide to bind.

Taken together, these findings show that the neuroendocrine system is indeed a major downstream target of hugin neurons and that this is achieved by a combination of synaptic and GPCR-mediated neuromodulatory transmission (Fig. 9E).

## Discussion

### Organizational principles of a peptidergic network

Almost all neurons in *Drosophila* are uniquely identifiable and stereotyped [63,64]. This enabled us to identify and reconstruct a set of 20 peptidergic neurons in an ssTEM volume spanning an entire larval CNS [27]. These neurons produce the neuropeptide hugin and have previously been grouped into four classes based on their projection targets (Fig. 2A) [20]. Our analysis allows detailed comparisons between neurons of the same class to address why the CNS sustains multiple copies of morphologically very similar neurons. We found that neurons of the same morphological class (a) were very similar with respect to the distribution of synaptic sites, (b) shared a large fraction of their pre- and postsynaptic partners and (c) in case of the interneuron classes (hugin-PC and hugin-VNC), neurons were reciprocally connected along their axons with other neurons of the same class. Similar features have been described for a population of neurons which produce crustacean cardioactive peptide (CCAP) in *Drosophila* [65]. The reciprocal connections as well as the overlap in synaptic partners indicate that the activity of neurons within each interneuron class is coordinately regulated and could help sustain persistent activity within the population. In the mammalian pyramidal network of the medial prefrontal cortex, reciprocal connectivity between neurons is thought to contribute to the networks robustness by synchronizing activity within subpopulations and to support persistent activity [66]. In the hypothalamus, interconnectivity and shared synaptic inputs, as described here for each hugin class, has been demonstrated for peptidergic neurons producing gonadotropin-releasing hormone (GnRH) and oxytocin [67,68]. Likewise, this is thought to synchronize neuronal activity and allow periodic bursting.

In addition, our results complement previous findings that specific phenotypes and functions can be assigned to certain classes of hugin neurons: hugin-VNC neurons were shown to increase locomotion motor rhythms but do not affect food intake whereas hugin-PC neurons modulate feeding behavior and are necessary for processing of bitter gustatory cues [19,40]. In accordance with this, we found that all hugin classes have their own unique sets of postsynaptic partners. This suggests that all hugin classes have specific, separable effects in a range of different contexts. One conceivable scenario would have been that each hugin class mediates specific aspects of an overarching ”hugin phenotype”. This would require that under physiological conditions all hugin classes are coordinately active. However, we did not find any evidence of such coordination on the level of synaptic connectivity. Instead, each hugin class forms an independent microcircuit with its own pre- and postsynaptic partners. This is in accordance with above-mentioned results on hugin-PC and hugin-VNC neurons. We thus predict that each class of hugin-producing neurons has a distinct context and function in which it is relevant for the organism. This makes hugin a valuable model system for studying how the same neuropeptide can serve multiple, unrelated functions. The first functional description of hugin was done in larval and adult *Drosophila* [18] while more recent publications have focused entirely on the larva [19,40]. One of the main reasons for this is the smaller behavioral repertoire of the larva: the lack of all but the most fundamental behaviors makes it well suited to address basic questions. Nevertheless, it stands to reason that elementary circuits should be conserved between larval and adult flies. To date, there is no systematic comparison of hugin across the life cycle of *Drosophila*. However, there is indication that hugin neurons retain their functionality from larva to the adult fly. First, morphology of hugin neurons remains virtually the same between larva and adults [18]. Second, hugin neurons seem to serve similar purposes in both stages: they acts as a brake on feeding behavior – likely as response to aversive sensory cues [19,40]. In larvae, artificial activation of this brake shuts down feeding [19]. In adults, removal of this break by ablation of hugin neurons leads to a facilitation (earlier onset) of feeding [18]. Such conserved function of neuropeptidergic function between larval and adult *Drosophila* has been observed only in a few cases. Prominent examples are short [69,70] and long neuropeptide F [71,72], both of which show strong similarities with mammalian NPY. The lack of other examples is not necessarily due to actual divergence of peptide function but rather due to the lack of data across both larva and adult. Given the wealth of existing data on hugin in larvae, it would be of great interest to investigate whether and to what extent the known features (connectivity, function, etc.) of the system are maintained throughout *Drosophila’s* life history.

### Parallel synaptic and neuromodulatory connections

A neural network is a highly dynamic structure and is subject to constant change, yet it is constrained by its connectivity and operates within the framework defined by the connections made between its neurons [73]. On one hand this connectivity is based on anatomical connections formed between members of the network, namely synapses and gap junctions. On the other hand, there are non-anatomical connections that do not require physical contact due to the signaling molecules, such as neuropeptides/-hormones, being able to travel considerable distances before binding their receptors [59]. Our current integrated analysis of the operational framework for a set of neurons genetically defined by the expression of a common neuropeptide, positions hugin-producing neurons as a novel component in the regulation of neuroendocrine activity and the integration of sensory inputs. We show that most hugin neurons receive chemosensory input in the subesophageal zone, the brainstem analog of *Drosophila* [19,24]. Of these, one class is embedded into a network whose downstream targets are median neurosecretory cells (mNSCs) of the pars intercerebralis, a region homologous to the mammalian hypothalamus [25]. We found that hugin neurons target mNSCs by two mechanisms. First, by classic synaptic transmission: Our data strongly suggest that acetylcholine (ACh) acts as transmitter at these synapses and subsets of mNSCs have been shown to express a muscarinic ACh receptor [58]. Second, by non-anatomical, neuromodulatory transmission using a peptide-receptor connection, as demonstrated by the expression of hugin G-Protein coupled receptor (GPCR) PK2-R1 (CG8784) in mNSCs. Strikingly, while PK2-R1 is expressed in all mNSCs, the hugin neurons are strongly synaptically connected to insulin-producing cells, but only weakly connected to DMS and DH44 neurons This mismatch in synaptic vs. peptide targets among the NSCs suggests an intricate influence of hugin-producing neurons on this neuroendocrine center. In favor of a complex regulation is that those mNSCs that are synaptically connected to hugin neurons also express a pyrokinin-1 receptor (PK1-R, CG9918) which, like PK2-R1, is related to mammalian neuromedinU receptors [74–76]. There is some evidence that PK1-R might additionally be activated by the hugin neuropeptide, which would add another regulatory layer [75]. The concept of multiple messenger molecules within a single neuron is well established and appears to be widespread among many organisms and neuron types [60,77–80]. For example, cholinergic transmission plays an important role in mediating the effect of NMU in mammals. This has been demonstrated in the context of anxiety but not yet for feeding behavior [81,82].

There are however only few examples of simultaneous employment of neuromodulation and fast synaptic transmission in which specific targets of both messengers have been investigated at single cell level. Prominent examples are AgRP neurons in the mammalian hypothalamus: these neurons employ neuropeptide Y, the eponymous agouty -related protein (AgRP) and the small molecule transmitter GABA to target pro-opiomelanocortin (POMC) neurons to control energy homeostasis [83]. Also reminiscent of our observations is the situation in the frog sympathetic ganglia, where preganglionic neurons use both ACh and a neuropeptide to target so-called C cells but only the neuropeptide additionally targets B cells. In both targets the neuropeptide elicits late, slow excitatory postsynaptic potentials [84]. It is conceivable that hugin-producing neurons act in a similar manner by exerting a slow, lasting neuromodulatory effect on all mNSCs and a fast, transient effect exclusively on synaptically connected mNSCs.

In addition to the different timescales that neuropeptides and small molecule transmitters operate on, they can also be employed under different circumstances. It is commonly thought that low frequency neuronal activity is sufficient to trigger fast transmission using small molecule transmitters, whereas slow transmission employing neuropeptides require high frequency activity [60]. Hugin-producing neurons could employ peptidergic transmission only as a result of strong excitatory (e.g. sensory) input. On the other hand there are cases in which base activity of neurons is already sufficient for graded neuropeptide release: *Aplysia* ARC motor neurons employ ACh as well as neuropeptides and ACh is released at lower firing rates than the neuropeptide. Nevertheless, peptide release already occurs at the lower end of the physiological activity of those neurons [85,86]. It remains to be seen how synaptic and peptidergic transmission in hugin neurons relate to each other.

The present study is one of very few detailed descriptions of differential targets of co-transmission and - to our knowledge - the first of its kind in *Drosophila*. We hope these findings in a genetically tractable organism will provide a basis for elucidating some of the intriguing modes of action of peptidergic neurons.

### Hugin as functional homolog of central neuromedinU

The mammalian homolog of hugin, neuromedinU (NMU), is found in the CNS as well as in the gastrointestinal tract [22]. Its two receptors, NMUR1 and NMUR2, show differential expression. NMUR2 is abundant in the brain and the spinal cord whereas NMUR1 is expressed in peripheral tissues, in particular in the gastrointestinal tract [87]. Both receptors mediate different effects of NMU. The peripheral NMUR1 is expressed in pancreatic islet *ft* cells in humans and allows NMU to potently suppress glucose-induced insulin secretion [74]. The same study also showed that Limostatin (Lst) is a functional homolog of this peripheral NMU in *Drosophila:* Lst is expressed by glucose-sensing, gut-associated endocrine cells and suppresses the secretion of insulin-like peptides. The second, centrally expressed NMU receptor, NMUR2, is necessary for the effect of NMU on food intake and physical activity [17,88]. In this context, NMU is well established as a factor in regulation of the hypothalamo-pituitary axis [31,32] and has a range of effects in the hypothalamus, the most important being the release of corticotropin-releasing hormone (CRH) [16,89]. We show that a subset of hugin-producing neurons targets the pars intercerebralis, the *Drosophila* homolog of the hypothalamus, in a similar fashion: neuroendocrine target cells in the pars intercerebralis produce a range of peptides, including diuretic hormone 44 which belongs to the insect CRH-like peptide family [90] (Fig. 10). Given these similarities, we argue that hugin is a functional homolog of central NMU just as Lst is a functional homolog of peripheral NMU. Demonstration that central NMU and hugin circuits share similar features beyond targeting neuroendocrine centers, e.g. the integration of chemosensory inputs, will require further studies on NMU regulation and connectivity.

**Figure 10.**
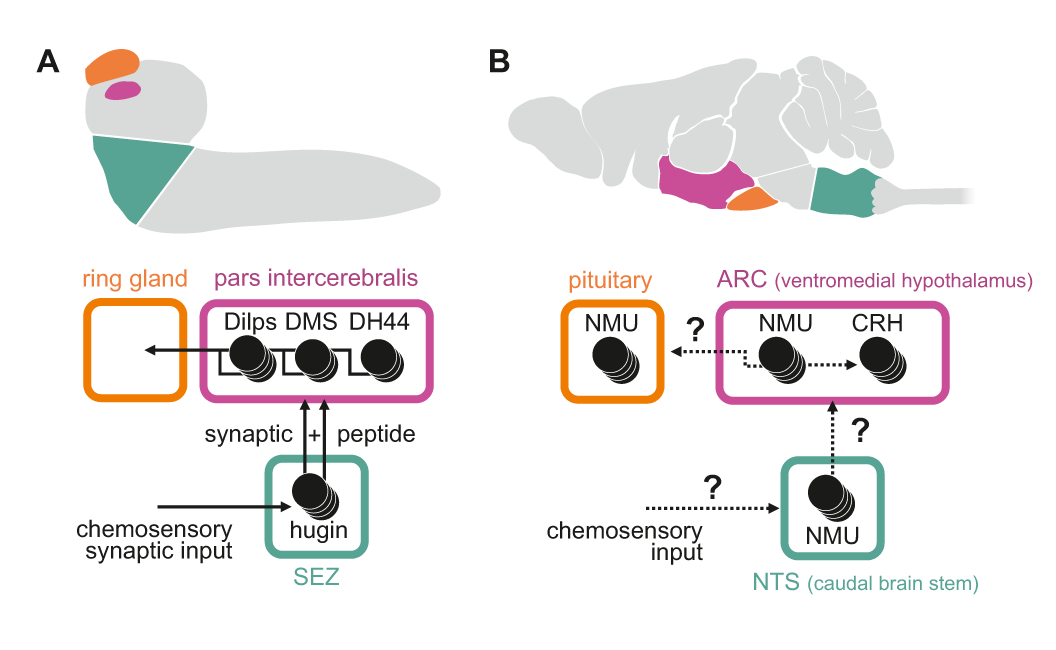
Summary of hugin connectivity and hypothetical implications for neuromedinU in mammals. **A**, Hugin neurons link chemosensory neurons that enter the subesophageal zone (SEZ) and neuroendocrine cells of the pars intercerebralis by synaptic as well as peptide-receptor connections. **B**, Distribution of NMU-positive neurons in mammals is much more complex, yet similar. The effect of neuromedinU (NMU) on feeding and physical activity originates in the arcuate nucleus (ARC) of the hypothalamus where it causes release of corticotropin-releasing hormone (CRH) which itself is a homolog of diuretic hormone 44 (DH44) in *Drosophila.* NMU-positive neurons have also been found in the nucleus of the solitary tract (NTS) a chemosensory center in the caudal brain stem. It remains to be seen if, similar to hugin neuron, NMU neurons serve as a link between chemosensory and neuroendocrine system.

Previous work on vertebrate and invertebrate neuroendocrine centers suggests that they evolved from a simple brain consisting of cells with dual sensory/neurosecretory properties, which later diversified into optimized single-function cells [35]. There is evidence that despite the increase in neuronal specialization and complexity, connections between sensory and endocrine centers have been conserved throughout evolution [34,36,37]. We argue that the connection between endocrine and chemosensory centers provided by hugin neurons represents such a conserved circuit that controls basic functions like feeding, locomotion, energy homeostasis and sex.

Indisputably, the NMU system in mammals is much more complex as NMU is found more widespread within the CNS and almost certainly involves a larger number of different neuron types. This complexity however only underlines the use of allegedly simple organisms such as *Drosophila* to generate a foundation to build upon. In summary, our findings should encourage research in other organisms, such as the involvement of NMU and NMU homologs in relaying chemosensory information onto endocrine systems.

## Acknowledgments

We thank Jan Veenstra and Liliane Schoofs for their gifts of antisera, and Hubert Amrein, Ron Tanimoto, Leslie Vosshall, Christian Jiinger, Barret Pfeiffer and Gerry Rubin for plasmids and fly lines. We thank SFB 645 and 704, DFG Cluster of Excellence ImmunoSensation and DFG grant PA 787 for financial support. We thank the Fly EM Project Team at HHMI Janelia for the gift of the EM volume, the HHMI visa office, and HHMI Janelia for funding. We also thank Lucia Torres, Gaia Tavosanis, Gaspar Jekely, Gregory Jefferis, Ingo Zinke, Scott Sternson, Christian Wegener, Volker Hartenstein and Nicholas Strausfeld for critical comments on earlier versions of this manuscript. The EM image data is available via the Open Connectome Project (http://www.openconnectomeproject.org).

## Materials and Methods

### Neuronal Reconstruction

Reconstructions were based on a ssTEM (serial section transmission electron microscope) data set comprising an entire central nervous system and the ring gland of a first-instar *Drosophila* larva. Generation of this data set was described previously [27]. Neurons’ skeletons were manually reconstructed using a modified version of CATMAID (http://www.catmaid.org) [91]. Hugin-PH (pharynx) neurons were first identified by reconstructing all axons in the prothoracic accessory nerve, through which these neurons exit the CNS towards the pharynx. Similarly, hugin-RG (ring gland) neurons were identified by reconstructing all neurosecretory cells that target the ring gland. To find the remaining hugin neurons, neighbors of already identified hugin neurons were reconstructed. Among those, the remaining hugin neurons were unambiguously identified based on previously described morphological properties such as projection targets, dendritic arborizations, relative position to each other and prominent landmarks like antennal lobes or nerves [20,92]. The mapped synaptic connections represent fast, chemical synapses matching previously described typical criteria: thick black active zones, pre- (e.g. T-bar, vesicles) and postsynaptic membrane specializations [38]. Hugin inputs and outputs were traced by following the pre- and postsynaptically connected neurites to the respective neurons’ somata or nerve entry sites in sensory axons. Subsequently, all sensory and endocrine neurons synaptically connected to hugin neurons were fully reconstructed. Interneurons were fully reconstructed if (a) homologous neurons were found in both hemispheres/-segments (did not apply to medially unpaired neurons) and (b) at least one of the paired neurons was connected by a minimum of 3 synapses to/from hugin neurons. Neurons that did not fit either criteria were not fully reconstructed and thus excluded from statistical analysis. This resulted in the reconstruction 177 synaptic partners that together covered 90%/96% of hugin neurons’ above threshold pre-/postsynaptic sites (Fig. 5 – figure supplement 1). The same parameters were applied to the reconstruction of synaptic partners of medial median neurosecretory cells (mNSCs). Morphological plots and example synapse’s volume reconstruction were generated using custom python scripts or scripts for Blender 3D (www.blender.org). The script for a CATMAID-Blender interface is on Github (https://github.com/schlegelp/CATMAID-to-Blender). See supplemental neuron atlas (Suppl. Files 1, 2) of all reconstructed neurons and their connectivity with hugin neurons.

### Localization of DCVs in respect to synaptic sites

Due to the neuronal reconstructions’ being skeletons instead of volumes, distances were measured from the center of each given dense core vesicle to the center of the closest presynaptic site along the skeleton’s arbors. DCVs within 3000 nm radius around the centers of neurons’ somata were excluded. Data was smoothed for graphical representation (Fig. 9D; bin size 50 nm).

### Normalized connectivity similarity score

To compare connectivity between neurons (Fig. 5B), we used a modified version of the similarity score described by Jarrell et al. [93]:

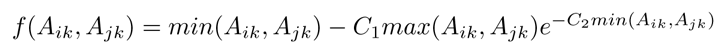

With the overall connectivity similarity score for vertices i and j in adjacency matrix A being the sum of *f* (*A*_*ik*_, *A*_*jk*_) over all connected partners k. *C*_1_ and *C*_2_ are variables that determine how similar two vertices have to be and how negatively a dissimilarity is punished. Values used were: *C*_1_ = 0.5 and *C*_2_ =1. To simplify graphical representation, we normalized the overall similarity score to the minimal (sum of –*C*_1_*max*(*A*_*ik*_, *A*_*jk*_) over all k) and maximal (sum of *max*(*A*_*ik*_, *Aj*_*k*_) over all k) achievable values, so that the similarity score remained between 0 and 1. Self-connections (*A*_*ii*_*A*_*jj*_) and *A*_*ij*_ connections were ignored.

### Synapse similarity score

To calculate similarity of synapse placement between two neurons, we calculated the synapse similarity score (Fig. 6):

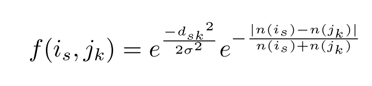

With the overall synapse similarity score for neurons i and j being the average of *f*(*i*_*s*_, *j*_*k*_) over all synapses s of i. Synapse k being the closest synapse of neuron j to synapses s [same sign (pre-/postsynapse) only]. *d*_*sk*_ being the linear distance between synapses s and k. Variable *σ* determines which distance between s and k is considered as close. *n*(*j*_*k*_) and *n*(*i*_*s*_) are defined as the number of synapses of neuron j/i that are within a radius ω of synapse k and s, respectively (same sign only). This ensures that in case of a strong disparity between *n*(*i*_*s*_) and *n*(*j*_*k*_), *f*(*i*_*s*_, *j*_*k*_) will be close to zero even if distance *d*_*sk*_ is very small. Values used: *σ* = *ω* = 2000 nm.

## Clustering

Clusters for dendrograms were created based on the mean distance between elements of each cluster using the average linkage clustering method. Clusters were formed at scores of 0.2 for synapse similarity score (Fig. 6B,E) and 0.4 for connectivity similarity score (Fig. 7D).

## Percentage of synaptic connections

Percentage of synaptic connections was calculated by counting the number of synapses that constitute connections between neuron A and a given set of pre- or postsynaptic partners (e.g. sensory neurons) divided by the total number of either incoming or outgoing synaptic connections of neuron A. For presynaptic sites, each postsynaptic neurite counted as a single synaptic connection.

## Statistics

Statistical analysis was performed using custom Python scripts; graphs were generated using Sigma Plot 12.0 (www.sigmaplot.com) and edited in Adobe Corel Draw X5 (www.corel.com).

## Generation of CG8784 promoter lines

The CG8784-GAL4::p65 construct (Fig. 8) was created using recombineering techniques [94] in P[acman] bacterial artificial chromosome (BAC) clone CH321-45L05 [95] (obtained from Children’s Hospital Oakland Research Institute, Oakland, CA), containing CG8784 within ≈80 kb of flanking genomic context. A generic landing-site vector was created by flanking the kanamycin-resistance/streptomycin-sensitivity marker in pSK+-rpsL-kana [96] (obtained from AddGene.org, plasmid #20871) with 5Ȉ and 3’ homology arms (containing GAL4 coding sequences and HSP70 terminator sequences, respectively) amplified from pBPGUw [97]. CG8784-specific homology arms were added to this cassette by PCR using the following primers (obtained as Ultramers from Integrated DNA Technologies, Inc., Coralville, Iowa; the lower-case portions are CG8784-specific targeting sequences, and the capitalized portions match the pBPGUw homology arms):

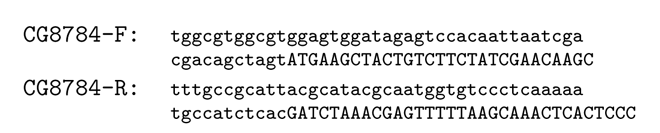

This cassette was recombined into the BAC, replacing the coding portion of the first coding exon, and then full-length GAL4::p65-HSP70 amplified from pBPGAL4.2::p65Uw [98] was recombined into the landing site in a second recombination. Introns and exons following the insertion site were retained in case they contain expression-regulatory sequences, although they are presumably no longer transcribed. Correct recombination was verified by sequencing the recombined regions, and the final BAC was integrated into the third-chromosome attP site VK00033 [99] by Rainbow Transgenic Flies, Inc. (Camarillo, CA).

The CG8784-6kb-GAL4 (Fig. 8 – figure supplement 1) was created using standard restriction-digestion/ligation techniques in pCaSpeR-AUG-Gal4-X vector [100]. An approximately 6-kb promoter fragment 5’ of the first coding exon was amplified using the following primers and inserted into a pCaSpeR vector (Addgene.org, plasmid #8378) containing a start codon (AUG) and the GAL4 gene.

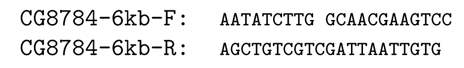

This construct was integrated into the genome via P-element insertion.

## Immunohistochemistry

For antibody stainings of CG8784-GAL4::p65, larvae expressing JFRC2-10xUAS-IVS-mCD8::GFP [98] driven by CG8784-GAL4::p65 were dissected in PBS. Brains were fixed in 4% formaldehyde in PBS for 1 h, rinsed, blocked in 5% normal goat serum, and incubated overnight at 4°C with primaries: sheep anti-GFP (AbD Serotec #4745-1051), 1:500; rabbit anti-DH44 [90] (gift of Jan Veenstra), 1:1000; rabbit anti-DILP2 [101] (gift of Jan Veenstra), 1:1000; 1:1000; and rabbit anti-DMS [56,102] (gift of Luc van den Bosch and Liliane Schoofs), 1:500. Tissues were rinsed and incubated overnight at 4°C in secondaries: Alexa Fluor 488 donkey anti-sheep (Jackson ImmunoResearch, #713-545-147) and rhodamine red-X donkey anti-rabbit (Jackson ImmunoResearch #711-296-152), both 1:500. Brains were rinsed and dehydrated through an ethanol-xylene series, mounted in DPX, and scanned on a Zeiss LSM 510 confocal microscope.

For antibody stainings of CG8784-6kb-GAL4, larvae expressing 10XUAS-mCD8::GFP (Bloomington, #32184) driven by CG8784-6kb-GAL4 were dissected in PBS. Brains were fixed in 4% paraformaldehyde for 30 minutes, rinsed, blocked in 5% normal goat serum, and incubated overnight at 4°C with primaries: goat anti-GFP-FITC (abcam, ab26662), 1:500; rabbit anti-DH44 [90] (gift of Jan Veenstra), 1:1000; guinea pig anti-Dilp2 [103] (Pankratz lab), 1:500 and rabbit anti-DMS [56,102] (gift of Luc van den Bosch and Liliane Schoofs), 1:500. Tissues were rinsed and incubated overnight at 4°C in secondaries: anti-rabbit Alexa Fluor 633 (Invitrogen, A-21070) and anti-guinea pig Alexa Fluor 568 (Invitrogen, A-11075), both 1:500. Brains were rinsed, mounted in Mowiol (Roth, 0713), and scanned on a Zeiss LSM 710 confocal microscope.

For antibody stainings against choline acetyltransferase (ChAT), larvae expressing a YFP-tagged halorhodopsin (UAS-eNpHR-YFP; Bloomington, #41753) driven by HugS3-GAL4 [18] as marker were prepared following the above protocol for CG8784-6kb-GAL4 stainings. Primary antibodies used: goat anti-GFP-FITC (abcam, ab26662), 1:500; mouse anti-ChAT (Developmental Studies Hybridoma Bank, ChAT4B1) [104], 1:1000. Secondary antibodies used: anti-mouse Alexa Fluor 633 (Invitrogen, A-21046). For investigation of ChAT promoter activity in hugin neurons, larvae expressing UAS-cd8a::mRFP (Bloomington, #27399) under the control of ChAT-GAL4 7.4kb (Bloomington, #6798) and YFP directly under the control of the hugin promoter (hug-YFP; [18]) were prepared following the above protocol for CG8784-6kb-GAL4 stainings. Primary antibodies used: goat anti-GFP-FITC (abcam, ab26662), 1:500; mouse anti-RFP (abcam, ab65856), 1:500. Secondary antibodies used: anti-mouse Alexa Fluor 633 (Invitrogen, A-21046).

For quantification of ChAT antibody signals/ChAT promoter activity, samples were scanned on a Zeiss LSM 710 confocal microscope using a 63X objective (Zeiss). Settings were kept the same over all scans. Regions of interest were placed through the center of each hugin neuron’s soma and the mean intensity was measured using ImageJ (https://imagej.nih.gov/ij/index.html) [105]. Hugin-PC and hugin-RG neurons were identified based on soma position and morphology. Hugin-VNC and hugin-PH could not be unambiguously discriminated as they were usually too tightly clustered. They were thus treated as a single group. For background normalization an approximately 10x10 *μm* rectangle from the center of the image stack was chosen.

## RNAi experiments

To investigate the role of acetylcholine as transmitter of hugin neurons, food intake and electrophysiological experiments were performed. Experimental procedures, materials and setups used in these assays been described extensively in Schoofs et al. (2014) [19]. The original RNAi data were used and expanded by ChAT RNAi experiments. The following GAL4 driver and UAS effector lines were used: HugS3-GAL4 [18], UAS-dTrpA1 (Bloomington, #26263), UAS-LacZRNAi (gift from M. Jünger), UAS-HugRNAi1A [19] and UAS-ChAT-RNAi (TriP.JF01877) (Bloomington, #25856) [43,47]. Controls consist of pooled data from wildtype (OregonR), w1118 and wildtype crossed with UAS-TrpA animals. There was no statistical difference between these control genotypes. For the food intake assay, third instar larvae were first washed and starved for 30mins on RT. They were then transferred on yeast paste colored with crimson red and allowed to feed for 20mins. Experiments were performed at 32°C for dTrpA-induced activation of hugin neurons and at 18°C as control condition. Afterwards larvae were photographed and the amount of food ingested was calculated as the area of the alimentary tract stained by the colored yeast divided by body surface area using ImageJ (https://imagej.nih.gov/ij/index.html) [105]. Data is represented as fold change between control condition (18°C) and dTrpA-induced activation (32°C) normalized to the control.

For the electrophysiological assay, semi intact preparations of third instar larvae were made in saline solution [106]. En passant extracellular recordings of the antennal nerve (AN) were performed following previously described protocol [19]. During the recordings, temperature of the CNS was alternated between 18°C (control condition) and 32°C (dTrpA activation). For analysis, fictive motor patterns of the pharyngeal pump (also: cibarial dilator musculature, CDM) were analyzed: fold change in cycle frequency between pairs of successive 18°C and 32°C sections of a recording was calculated.

### Pharmacological experiments and calcium imaging

Hugin-derived pyrokinin 2 (hug-PK2) was synthesized by Iris Biotech (Marktredwitz, Germany) using the amino acid sequence SVPFKPRL-NH2. The C-terminus was amidated. The effect of hug-PK2 on calcium activity in median neurosecretory cells (mNSCs) was investigated using the calcium integrator CaMPARI [107]. To drive expression of CaMPARI in mNSCs, CG8784-6kb-GAL4 flies were crossed to UAS-CaMPARI (Bloomington #58761). Larval brains were dissected and placed in saline solution [106] containing either no, 100*nM*, 1*μM* or 10*μM* hug-PK2. After 1min of incubation, 405nm photoconversion light was applied for 15s. Afterwards brains were placed on a poly-l-lysine-coated (Sigma-Aldrich, P8920) cover slide and scanned using a Zeiss LSM 780 confocal microscope. Settings were kept the same over all scans. Calcium activity was calculated as the ratio of the fluorescence of photoconverted (red) to unconverted (green) CaMPARI using ImageJ.

## Figure Supplements

**Figure 2. – figure supplement 1:**
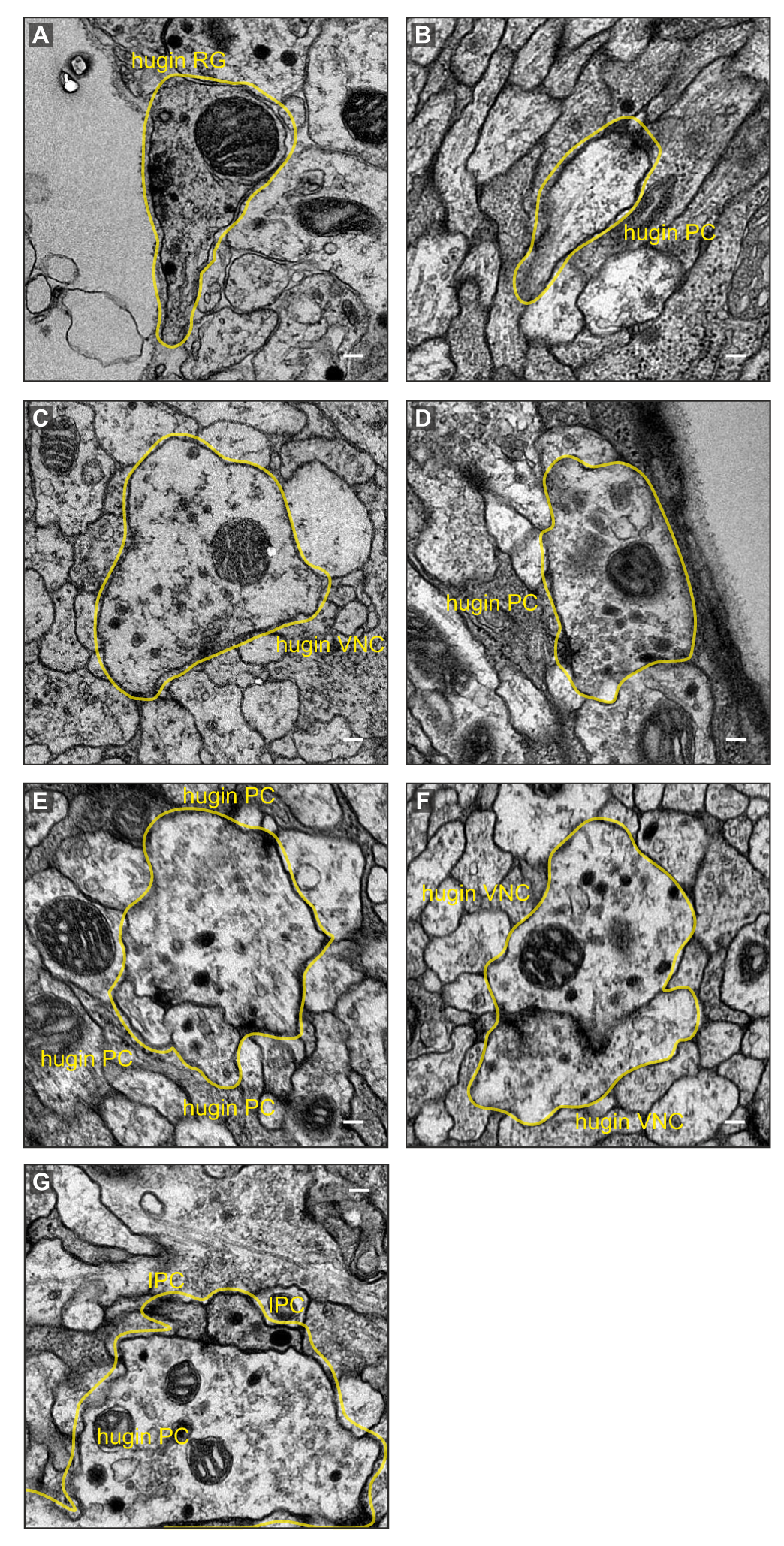
Examples of synaptic sites in the ssTEM volume. **A**, Presynaptic density of a hugin-RG neuron bordering haemal space within the ring gland. **B**,**C**, Examples of presynaptic sites with small clear core vesicles for a hugin-PC and hugin-VNC neuron. **D**, Example of a presynaptic site with close-by dense core vesicles. **E**, **F**, Examples of synaptic connections between hugin neurons. **G**, Examples of synaptic connections from hugin-PC neurons onto insulin-producing cells (IPCs). Scale bars represent 100nm.

**Figure 5. – figure supplement 1:**
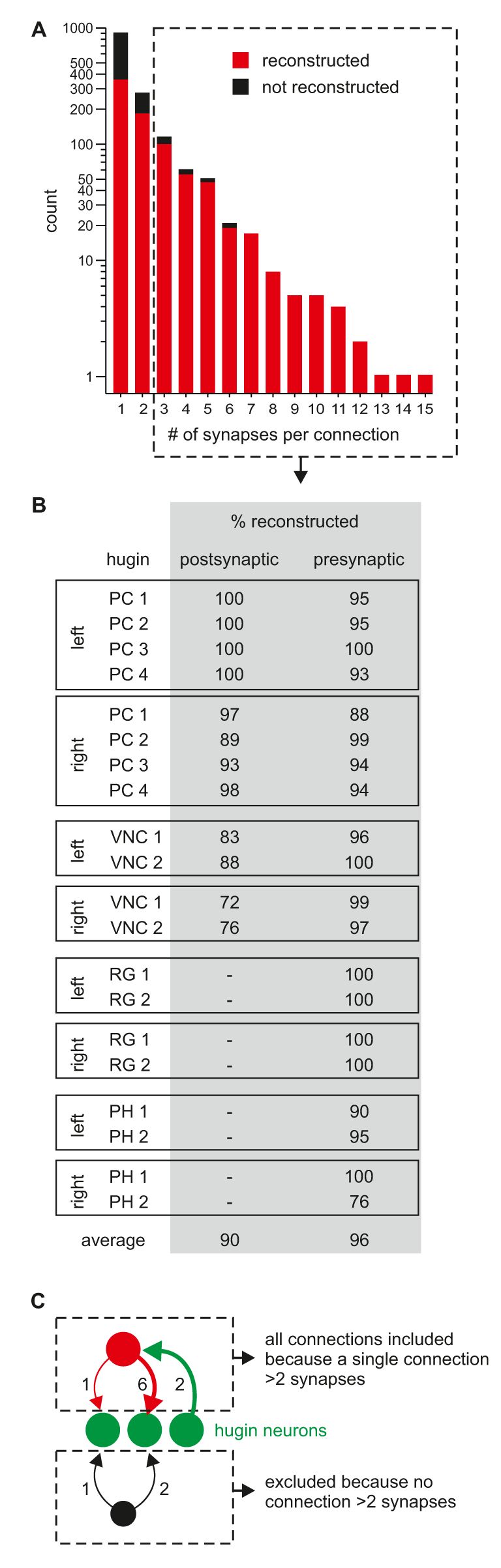
Neurons connected by more than two synapses to hugin neurons were reliably reconstructed. **A**, Distribution of synaptic connections to/from hugin neurons. X-axis gives number of synapse per connection and y-axis occurrence. Reconstruction of the pre- or postsynaptic neuron was attempted for every hugin synapse. Red fraction was completely reconstructed, black fraction was not successfully reconstructed due to either errors/ambiguity in the ssTEM data set (e.g. missing sections) or failure to find a matching pair of neurons in both hemisegments. The fraction of unaccounted synapses strongly decreases from 2 to 3 synapses per connection. We therefore subsequently applied a threshold of at least a single more-than-two-synapses connection (did not apply to sensory neurons). **B**, Completeness of reconstruction of pre- and postsynaptic partners for each hugin neuron. Values in the table give the percentages of pre- and postsynaptic sites that connect to an above-threshold partner of hugin neurons which was successfully reconstructed.

**Figure 6. – figure supplement 1:**
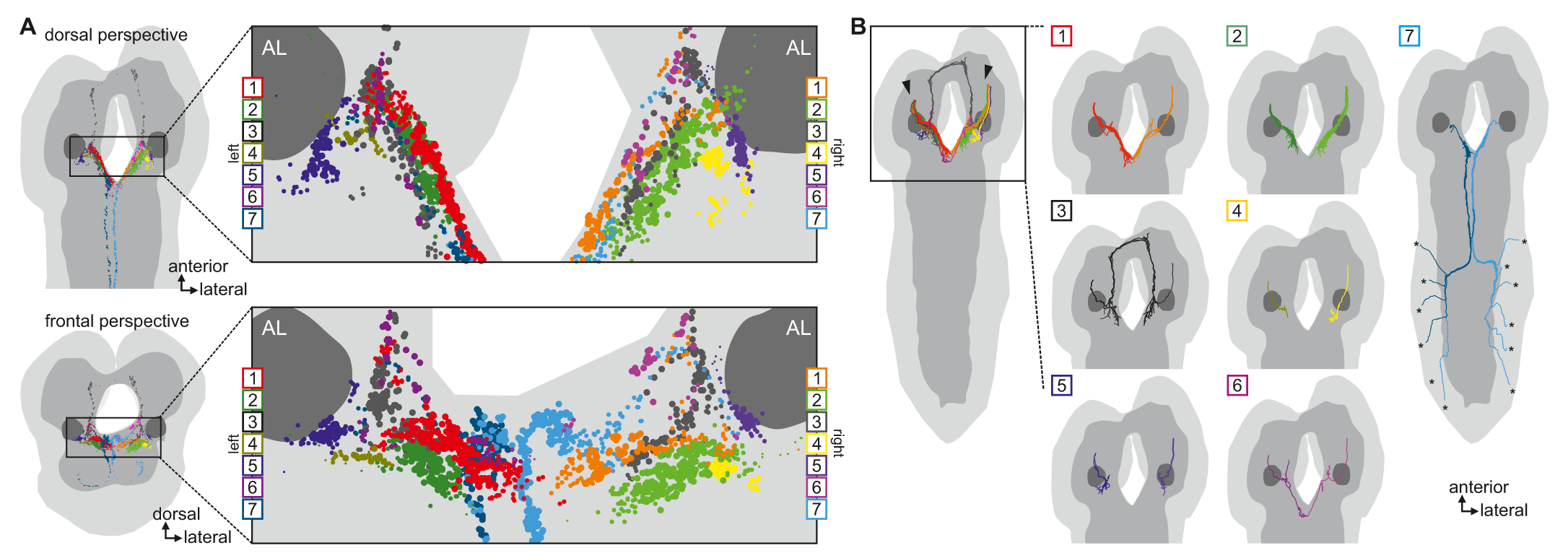
Clustered synapses of sensory inputs to hugin neurons cover discrete parts of the subesophageal zone. Distribution of synaptic sites of afferent neurons as clustered in Fig 6. Each dot indicates a synaptic site. Dot size decreases with distance to its cluster’s center. Note, that the sensory neurons presynaptic to hugin neurons innervate areas medial and ventral to the antennal lobes. See also video 3.

**Figure 8. – figure supplement 1:**
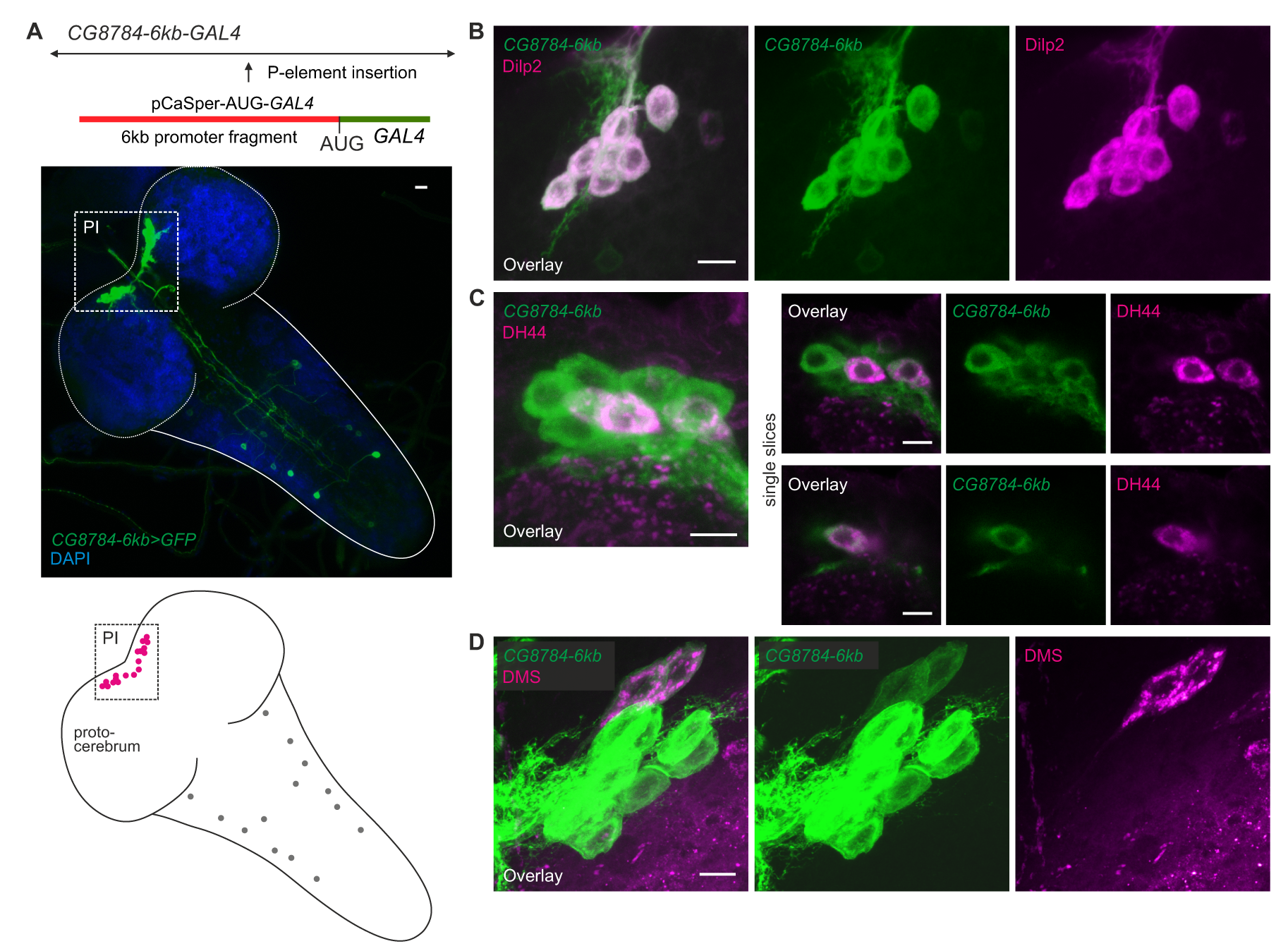
Hugin receptor line CG8784-6kb-GAL4 drives expression in median neurosecretory cells (mNSCs) of the pars intercerebralis (PI) similar to CG8784-GAL4:p65. **A**, Immunolabeling and semi-schematic representation of hugin receptor CG8784-6kb-GAL4 expression pattern. In comparison with the CG8784-GAL4:p65 BAC line (Fig. 8), this line shows a more restricted expression. However, the same prominent cluster of neurons (magenta) of the PI is labeled but only few additional cells in the ventral nerve cord (grey circles). **B**-**D**, Double staining of CG8784-6kb-GAL4 driving GFP expression suggests expression of CG8784 in (B) *Drosophila* insulin-like peptide- (Dilp2), (C) diuretic hormone 44- (DH44) and (D) Dromyosuppressin-producing (DMS) mNSCs of the PI. Scale bars represent 10 μm (A) and 5 μm (B-D).

**Figure 8. – figure supplement 2:**
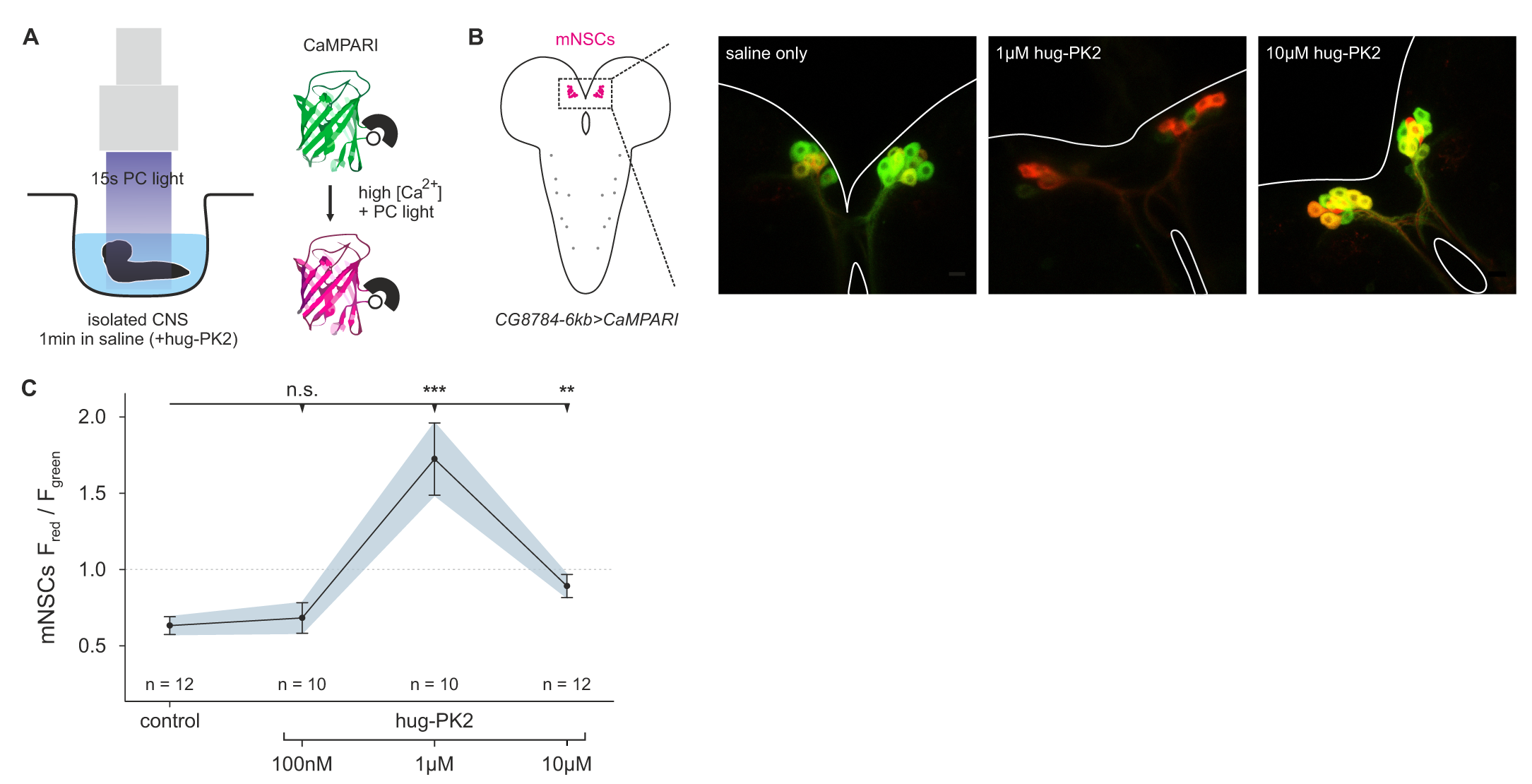
Hugin neuropeptide increases calcium activity in median neurosecretory cells (mNSCs). **A**, Experimental setup. Isolated central nervous systems (CNS) expressing the calcium integrator CaMPARI were dissected and placed in saline solution or saline solution + hug-PK2 (hugin-derived pyrokinin2). After 1min incubation 405nm photoconversion (PC) light was applied for 15s. Afterwards brains were scanned and ratio of converted (red) to unconverted (green) CaMPARI was analyzed. **B**, Exemplary scans of mNSCs after incubation and photoconversion in saline control or saline + hug-PK2. CG8784-6kb-GAL4 was used to drive expression of UAS-CaMPARI in mNSCs. **C**, Quantification of calcium activity in mNSCs after incubation with hug-PK2 as measured by ratio of red to green fluorescence (F_*red*_/F_*green*_). Incubation with 1*μM* or 10*μM* hug-PK2 significantly increases calcium activity.

## Supporting Information

**Supplementary File 1: PDF Neuron Atlas - Morphology and connectivity of reconstructed neurons.** Reconstructions of (A) hugin-PC, (B) hugin-VNC, (C) hugin-RG, (D) hugin-PH neurons, (E) insulin-producing cells (IPCs), (F) DH44-producing cells, (G) DMS-producing cells, (H) antennal nerve (AN) sensory neurons, (I) abdominal nerve sensory neurons, (J) paired interneurons and (K) unpaired medial interneurons. A dorsal view of each cell is shown on the left, and a frontal view on the right. Neuron ids (e.g. #123456) are provided to allow comparison between PDF and Blender atlas. Outline of the nervous system and the ring gland are shown in grey and dark grey, respectively. Number in the table is the number of synapses of given neurons onto (left) and from (right) the hugin neuron indicated in the row. Neurons are displayed as corresponding pairs of the left/right hemisegment with the exception of sensory neurons and unpaired medial interneurons.

**Supplementary File 2: Blender 3D Neuron Atlas – Morphology of reconstructed neurons as Blender file.** To view, please download Blender (www.blender.org). Reconstructed neurons are sorted into layers: hugin neurons (1), mNSCs (2), sensory neurons (3), interneurons (4) and mesh of the larval brain (5, hidden by default). Neuron names contain id (e.g. #123456) to allow comparison between Blender and PDF atlas. Neurons have been resampled by a factor of 4 to reduce vertex count. 1nm = 0.0001 Blender units.

**Video 1: Morphology of hugin-producing neurons.** Video shows morphology of hugin-producing neurons as well as distribution of their presynaptic and postsynaptic sites. Hugin interneurons (PC and VNC) have mixed input-output compartments whereas efferent hugin neurons (PH and RG) show almost exclusively postsynaptic sites within the CNS. Outlines of the CNS including the ring gland are shown in white.

**Video 2: Each class of hugin neurons connects to unique sets of synaptic partners.** Video shows all reconstructed presynaptic and postsynaptic partners of hugin neurons (see Fig. 5C). Neurons are colored by total number of synapses to/from given hugin class. Each hugin class forms distinct microcircuits with little to no overlap with those of other classes.

**Video 3: Clusters of chemosensory neurons cover distinct areas of the subesophageal zone (SEZ).** Video shows morphology and presynaptic sites of sensory inputs to hugin neurons. Neurons are clustered based on a synapse similarity score (see Fig. 6C). Each sphere indicates a presynaptic site. Sphere size increases with the number of postsynaptically connected neurons.

